# A Robust Genome-Wide Association Study Uncovers Signature Genetic Alterations among *Pseudomonas aeruginosa* Cystic Fibrosis Isolates

**DOI:** 10.1101/2020.12.02.407528

**Authors:** Wontae Hwang, Ji Hyun Yong, Kyung Bae Min, Kang-Mu Lee, Sang Sun Yoon

## Abstract

*Pseudomonas aeruginosa* (PA) is an opportunistic pathogen that causes diverse human infections such as chronic airway infection in cystic fibrosis (CF) patients. Although many sequenced genomes are available, a comprehensive comparison between genomes of CF versus non-CF PA isolates remains yet to be conducted. In order to gain a deeper understanding into the PA adaptation in the CF airway, we performed a Genome-Wide Association Study (GWAS) using a total of 1,001 PA genomes. Genetic variations uniquely identified among CF isolates were categorized into (i) alterations in protein-coding regions either large- or small-scale and (ii) polymorphic variations in intergenic regions. We introduced each CF-specific genetic alteration into the genome of PAO1, a prototype PA strain and experimentally validated their outcomes. Loci readily mutated among CF isolates include genes encoding a probable sulphatase and a probable TonB-dependent receptor (PA2332~PA2336), L-cysteine transporter (YecS, PA0313) and a probable transcriptional regulator (PA5438). A promoter region of heme/hemoglobin uptake outer membrane receptor (PhuR, PA4710) was similarly identified as meaningfully different between the CF and non-CF isolate groups. Our analysis, the first of its kind, highlights how PA evolves its genome to persist and survive within the context of chronic CF infection.

## Introduction

*Pseudomonas aeruginosa* (PA) is a gram-negative bacterium commonly found in various places, such as soil and water, but it is also found in immunocompromised patients as an opportunistic pathogen [1]. PA can cause not only acute syndromes like pneumonia and bloodstream infection, but also chronic airway infections in cystic fibrosis (CF) patients. CF is a well-known genetic disorder caused by a mutated CFTR (cystic fibrosis transmembrane conductance regulator) protein. CFTR disruption alters the lung condition such that the increasingly dehydrated viscous mucus layer provides a favourable habitat to several pathogens, such as PA, over an extended period of time [2].

Evolutionary versatility of PA due to its large genome containing numerous regulatory genes gives the bacterium an advantage in adapting to prolonged infections [3]. For this reason, there are many studies that aspired to understand how PA adjusts and responds to the harsh CF lung environment. In order to gain insights on a genetic level, a small-scale mutation tracking analysis of PA isolates from one CF patient was performed by whole genome sequencing [4]. On a transcriptomic level, PA grown exclusively in CF sputum as the sole energy source presented altered expression of genes encoding functions of amino acid biosynthesis and degradation, and quinolone signalling [5, 6]. It was reported that mutations in *lasR* gene, which encodes an important quorum sensing regulator, are frequently detected among CF isolates [7, 8], although the significance of this mutation is debatable. Not surprisingly, mutations leading to increased antibiotic tolerance have been revealed in PA inhabiting the CF airway [9–11]. Furthermore, PA clone types vary across different CF patients, depending on the sputum composition [12–15] and even within an individual patient over time [16, 17]. Together, these results clearly suggest that PA is an adaptable organism that responds flexibly to changing environments.

Most of the genetic level studies conducted so far, however, mainly focused on CF isolates, but not non-CF isolates. For this reason, genetic variations revealed from such studies cannot be ascertained whether or not they are responsible for PA changes specifically induced by the CF environment. In the present study, we compared 1,001 PA genomes derived either from CF or non-CF clinical isolate. Genome-Wide Association Studies (GWASs) have proved useful in uncovering causal relationships between genetic variations and disease phenotypes in human populations [18]. According to the GWAS catalog [19], > 4,000 human GWASs have been conducted. In contrast to human genomes, bacterial genomes are variable even within the same species in terms of size, gene repertoire, and gene arrangement [20]. Due to this inherent feature of bacterial genomic plasticity, GWAS of bacterial populations has never been as active as for humans [21, 22]. In this work, we selected PA isolate genomes of known origins and performed a GWAS based on k-mer counting [22–24], a modified method permitting association mapping inside the intergenic regions and outside the PAO1 genome as well. Results provided here expand our current understanding of how gene-level changes correlate with mechanisms of PA adaptation to the CF lung environment.

## Results and Discussion

### Genome-wide association study (GWAS) of PA isolated from CF and non-CF patients

#### A. Genome selection and phylogenetic tree construction

A lot of previous genetic level studies conducted with the aim to better understand chronic PA infection in CF patients had targeted isolates from varying numbers of patients [14, 16]. While such analyses have contributed many meaningful insights, they focused only on CF isolates, and isolates from non-CF individuals were excluded from analysis. Thus, it is ambiguous whether genetic variations highlighted in these studies are important in the context of CF-specific adaptation or not. In this study, we downloaded 2,167 PA genomes from Pseudomonas genome database [25]. This selection contained genomes of isolates from both CF and non-CF individuals, and downstream analyses were performed to find genetic variants specifically found in either CF or non-CF isolates. After removing genomes of unknown origins, a phylogenetic tree was constructed with 1,001 genomes of known origins using RapidNJ [26], a phylogenetic tree construction tool employing the neighbour-joining method (Fig. S1A). In order to reduce the tree size, Treemmer [27] was used to exclude very similar genomes in clades. As a result, the tree originally constructed from 1,001 genomes was effectively trimmed down to contain 636 genomes which maintained 99.8 % diversity of the original selection (supplementary data 1) and the overall structure of the phylogenetic tree. In Fig. S1, black and red leaves of the phylogenetic tree indicate non-CF and CF isolates, respectively. As already known from a previous study by Ozer *et al.*, [28], most of the CF isolates belong to the main group (blue dotted line) which includes the majority of genomes, and we refer to the other genome group as the sub-group (red dotted line).

#### B. Pyseer results based on 31mer counting

Total 636 genomes consisting of 206 genomes from CF and 430 genomes from non-CF were used to perform Pyseer [29], a bioinformatic tool used for GWAS, to find specific variants that may be important for PA adaptation in CF airways. In our study, k-mer based GWAS was implemented because this method provides us more information regarding intergenic regions. We selected k-mer whose length is 31 base pairs (31mer), a measure that is not too sensitive or too specific, and calculated total 31mers from all genomes by Fsm-lite [30]. After Pyseer analysis with total 31mers as input, lrt-pvalues, which considers the population structure of all genomes, were calculated for each 31mer. All 31mers with bad-chi values or lrt-pvalues above 2.8e-08, the significance threshold in this analysis, were filtered out. The remaining 31mers were sorted as significantly different 31mers between CF and non-CF groups. Subsequently, these significant 31mers were aligned to PAO1 genome, the representative genome of the non-CF group, and the results were visualized by Phandango [31] (Fig. 1A). In Fig. 1A, asterisk regions, where lots of candidate 31mers aligned to, are ribosomal DNA (rDNA) sequences and the four peaks represent the four identical copies of rDNA sequences in the PAO1 genome. A greater number of CF isolates did not seem to have 16S and 23S rDNA sequences compared with the non-CF group. But genome sequencing of several CF isolates identified that there are rDNA sequences normally (data not shown). Despite the occurrence of such error in rDNA sequence recognition in computer based analysis, we included these genomes in the input database, because the number of annotated genes of those genomes as predicted by Prokka [32] were not significantly different from the rest of the genomes. While a lot of statistically meaningful 31mers were distributed across diverse regions of the PAO1 genome, one spot had a prominent number of 31mers aligned (Fig. 1A), and we refer to this region as the hot spot (hs). After de-novo assembling the total 31mers aligned to the PAO1 genome into contigs by Trinity [33], we found that 29 and 109 contigs were respectively bound to non-coding regions and genes capable of translation (except untranslated genes such as rDNA sequences due to the aforementioned issue).

**Figure 1.**
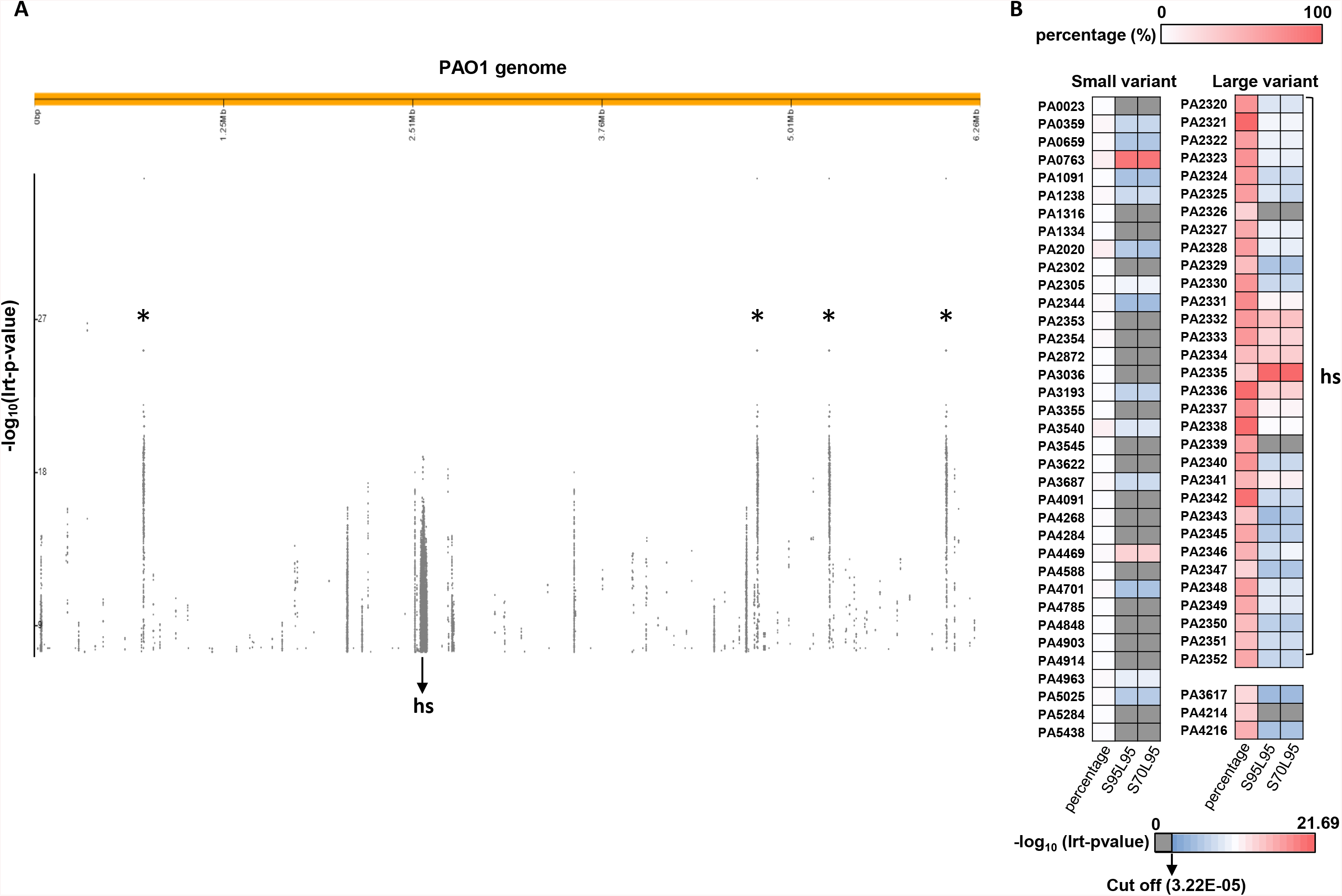
Visualization of significantly different regions and genes between CF and non-CF groups. **(A)** Significantly different 31mers between CF and non-CF groups were aligned to PAO1 whole genome and visualized by Phandango. Regions marked by asterisks are ribosomal DNA sequences and “hs” marks the hotspot region where a large number of candidate 31mers aligned to. The higher up a spot representing a 31mer is located, the more significant was its lrt-pvalue. **(B)** Various candidate proteins from the PAO1 genome are presented. Candidates were classified as a large variant, if the percentage of variable residues exceed 20% of the protein length, and if not, as a small variant. The leftmost column of boxes in each group indicates the percentage of the protein length the variable residues within the region makes up, as per the colour index above the heatmap. Lrt-pvalues calculated for clustering performed under two conditions (S95L95: 95% similarity and 95% length coverage; S70L95: 70% similarity and 95% length coverage) are described in the second and third columns as per the colour index beneath the heatmap. Candidates with gray lrt-pvalue boxes were determined to be insignificantly different between CF and non-CF groups. “hs” indicates candidate proteins located within the hotspot region in Fig. 1A.

#### C. Amino acid level examination of significant variants

In order to examine the significantly different variants on the amino acid level, we used the locus information of 109 contigs binding to translated coding regions, and converted the nucleotide sequence information into amino acid sequences. We then ran Pyseer again with the amino acid residue information as input. In a similar manner to sorting the previous Pyseer analysis results, we excluded amino acid residues with bad-chi values or lrt-pvalues above 4.01E-06, the cutoff value used in this analysis. As such, final amino acid residues were chosen. Detection of changes in the amino acid sequences was based on proteins of PAO1. Fig. 1B presents proteins that had at least one amino acid residue significantly different between CF and non-CF group, and was divided into small variant and large variant groups. A protein was categorised as a small variant if variable residues within that protein comprise less than 20% of protein length, and if not, the protein was classified as a large variant. Hence most small variants have their leftmost boxes coloured pale, which corresponds to less than 10% as described by the colour index above the heatmap. In contrast, large variants have their leftmost boxes coloured deeper red, as the percentage of variable residues in those protein exceed 20%.

Next, we set out to determine whether mutations in the small variant group lead to frameshifts and if the candidate proteins in the large variant group are indeed related to gene deletion. To this end, we first clustered homologues of each protein from entire genomes, based on the PAO1 genome using Blastclust [34], and various similarity and length coverage options were implemented (supplementary data 2). These clusters were used as input to run Pyseer. After removing clusters with lrt-pvalues larger than cutoff (3.22E-05) and bad-chi values, the remaining candidate protein clusters and their corresponding lrt-pvalues at two conditions (95% similarity and 95% length coverage; 70% similarity and 95% length coverage) are depicted in the second and third columns (Fig. 1B). The deeper the red colour of the boxes are in these two columns, the more significant difference is suggested between the CF and non-CF groups with regard to the protein cluster. Gray-coloured boxes in the second and third columns indicate that no significant difference between CF and non-CF isolates was observed in whole protein sequences. Detailed information about amino acid residues and clustering was provided in supplementary data 2 and supplementary data 3.

### Large-scale alterations in protein-coding regions: PA2332 ~ PA2336

In the large variant group, most candidate proteins were included in hs region, except for PA3617, PA4214, and PA4216 (Fig. 1B). In contrast to the small variant group, a greater number of large variants presented significant associations with either disease statuses (CF or non-CF), as represented by the fewer numbers of gray boxes in columns 2 and 3 (Fig. 1B). Among prominent genes with significantly small lrt-pvalues (from PA2332 to PA2336), PA2335, a probable TonB-dependent receptor, was detected with the smallest lrt-pvalue for its clustering result.

Total amino acid residues of PA2335 are represented in Fig. 2A and each amino acid locus is represented by the individual squares. Colours of the squares indicate the lrt-pvalue assigned to that amino acid residue. Based on the lrt-pvalues, correlations of PA2335 alteration with disease statuses (CF or non-CF) were significantly different, as determined by the clustering analysis performed with Pyseer (Fig. 2B). Unlike the lrt-pvalues, taking the population structure into calculation, filter-pvalues calculated for the entire genomes were very large, as they do not consider the population structure (Fig. 2B). These two types of p-values initially seem to suggest opposite conclusions about the importance of PA2335 alteration in PA adaptation to CF airway, and the importance of taking the population structure into consideration is portrayed in Fig. 2C. The phylogenetic tree is drawn using the genomes of PA isolates, and the three rows below describe the associated condition (CF or non-CF), and whether the isolate’s homologue clustered with PA2335 at given clustering options (S95L95; S70L95), in that order. Ideally, in order to support the notion that PA2335 alteration is critical in CF adaptation, three red lines would align at high frequency across the entire selection of genomes. Our results show that this is true, except for region **a**(Fig. 2C). This region corresponds to the entire sub-group in the unrooted phylogenetic tree (Fig. S1B, red dotted line). PA2335 alteration is very frequent in PA that belongs to region **a**, and does not seem to have any correlation with the CF isolates of that region. But CF isolates from outside region **a** show significantly high correlation with PA2335 alteration, as reflected in the small lrt-pvalues. PA2333, PA2334 and PA2336 present identical patterns as PA2335, whereas PA2332 presented both significantly small lrt-pvalue and filter-pvalue (data not shown). PA2332 alteration seems to be meaningfully associated with CF condition, across the entire population of isolates (supplementary data 2).

**Figure 2.**
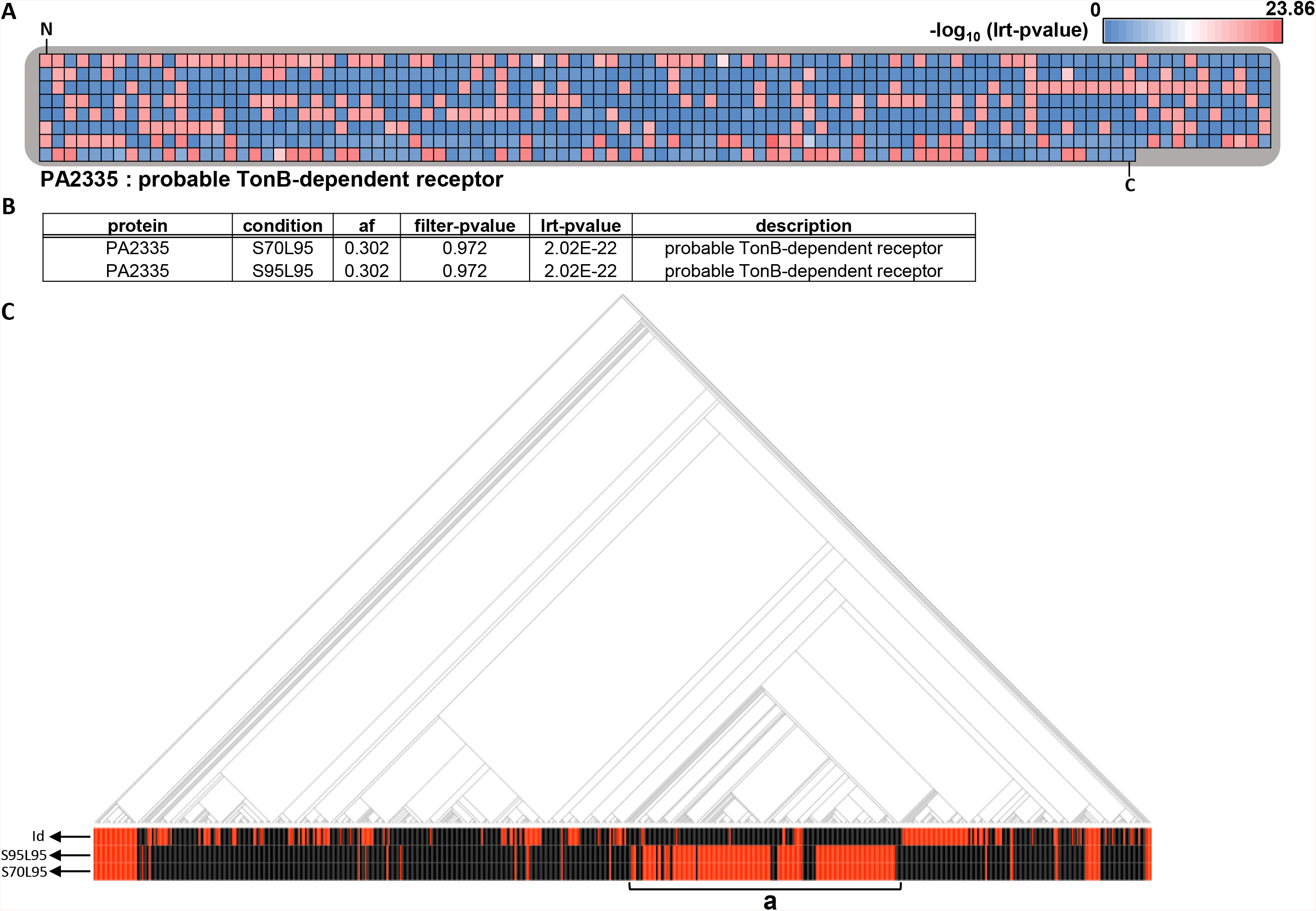
PA2335, candidate with the lowest lrt-pvalue in the large variant group. **(A)** Total amino acid residues of PA2335 are presented and each amino acid locus is represented by individual squares. Colour of each square indicates the lrt-pvalue assigned to that amino acid residue based on Pyseer results and colours were assigned according to the index above the heatmap, which was generated based on the negative logs of the maximum and minimum lrt-pvalues from Pyseer result of amino acid locus. “N” and “C” mark the N- and C-terminus of PA2335, respectively. **(B)** Pyseer result of the PA2335 cluster is shown. Different conditions used in clustering and the frequency of the PA2335 large alteration are described in “condition” and “af” columns. Population structure is not considered in calculation of filter-pvalues, whereas it is for calculation of lrt-pvalues. **(C)** Phylogenetic tree of 636 genomes and their associated disease status are shown. Red and black lines in the “id” row indicate genomes isolated from CF and non-CF patients, respectively. Black lines in the second and third rows indicate that the homologue of PA2335 clustered with the PAO1 protein at given clustering options (S95L95:S70L95), whereas red lines indicate that the homologue did not cluster with the PAO1 protein. Sub-group in **Fig. S1** is marked as **a**.

In a study by Ozer et al., (2019), they found lineage marker genes between main-group and sub-group (Fig. S1) whose distribution is skewed towards either group. For instance, *exoS* and *exoU* were mainly detected in the main-group and sub-group, respectively [28]. Similarly, PA2333~PA2336 qualify as a lineage marker, because most genomes in the sub-group do not seem to contain these genes unlike the main-group (Fig. 2C). Furthermore, we anticipate that there exists some difference in the genetic elements of isolates in the two groups, that contributes to the significance of alteration of PA2333~PA2336 in CF adaptation, whereas PA2332 alone is sufficient to establish significant effects regardless of the presence of such elements.

We excluded the large variant candidates (PA2326, PA2339 and PA4214), of which homologues presented no difference in clustering between CF and non-CF groups (Fig. 1B), and the remaining candidates were functionally annotated by Blastkoala [35]. KEGG ontology (KO), definition and pathway information of those candidates are listed in Table 1 [36]. Several transporters, pentose phosphate pathway enzymes and gene related to phenazine biosynthesis are included. Detailed information about whether each alteration is predominantly observed in the CF or non-CF groups is included in supplementary data 3.

**Table 1.**
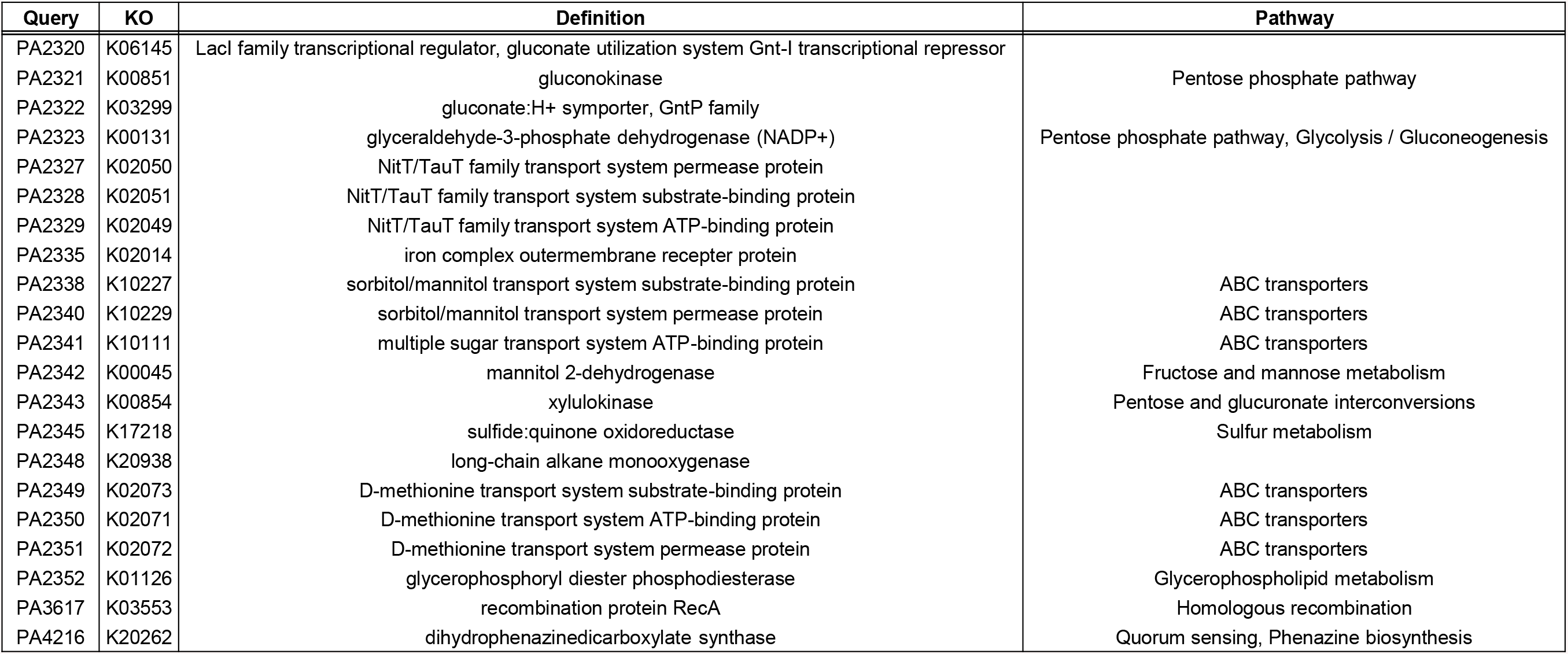
KEGG annotation of candidate genes in the large variant group. All large variants except for PA2326, PA2339 and PA4214 (with gray-coloured lrt-pvalue boxes in Fig. 1B) were annotated by Blastkoala. Proteins with KEGG information are listed in the “Query” column, and KEGG ontology (KO), definition and related pathway information are shown.

### Small-scale alterations in protein-coding regions

In our study, we focused more on the small variants that differ in presence between CF and non-CF isolates, as acquirement of small mutations may be more cost-effective compared to large indel mutations. Based on lrt-pvalues, top 20 amino acid residues of the small variant group whose 31mers aligned to the PAO1 genome are listed in Table 2. For instance, MucA (PA0763) is classified as a small variant (Fig. 1B), meaning that a small number of amino acid residues is altered compared to the reference PAO1 MucA protein. Clustering results of MucA homologues (at S95L95 and S70L95) presented significant lrt-pvalues, suggesting that variants of MucA implicate critical mutations rather than simple missense mutations. To test this, we compared the homologue sequences of MucA from each genome with the representative PAO1 MucA protein sequence. Interestingly, most small variants generated a premature stop codon in the *mucA* gene. Variants of MucA here served as positive control, as it is well known that mutations in the *mucA* gene are frequently found in CF isolates and that they contribute to the mucoid phenotype of the isolates [37–40].

**Table 2.** Top 20 amino acid residues of the small variant group whose 31mers aligned to the PAO1 genome. Top 20 amino acid residues according to Pyseer results whose 31mers aligned to the PAO1 genome are presented. The reference protein corresponding to each residue is presented in the “protein” column, and the “locus” column contains locus of the mutation and amino acid residue of the reference protein where the mutation is detected. Multiple types of mutation at each locus of the reference protein may be present and detailed information is provided in supplementary data 3.

| protein | locus | af | filter-pvalue | lrt-pvalue | description |
| --- | --- | --- | --- | --- | --- |
| PA2020 | 46G | 0.159 | 4.02E-17 | 1.65E-20 | MexZ |
| PA0763 | 124I | 0.119 | 1.72E-28 | 2.03E-17 | anti-sigma factor MucA |
| PA0763 | 125A | 0.119 | 1.72E-28 | 2.03E-17 | anti-sigma factor MucA |
| PA0763 | 127P | 0.119 | 1.72E-28 | 5.00E-17 | anti-sigma factor MucA |
| PA0763 | 123Q | 0.118 | 6.10E-28 | 5.88E-17 | anti-sigma factor MucA |
| PA5438 | 272S | 0.075 | 7.03E-20 | 9.92E-17 | probable transcriptional regulator |
| PA0763 | 126L | 0.119 | 3.07E-27 | 1.46E-16 | anti-sigma factor MucA |
| PA2020 | 45R | 0.118 | 8.88E-07 | 3.86E-15 | MexZ |
| PA5438 | 273L | 0.079 | 1.20E-19 | 4.41E-15 | probable transcriptional regulator |
| PA4469 | 146L | 0.104 | 1.41E-25 | 7.93E-15 | hypothetical protein |
| PA4469 | 148Y | 0.104 | 1.41E-25 | 7.93E-15 | hypothetical protein |
| PA4469 | 147R | 0.105 | 4.07E-26 | 1.39E-14 | hypothetical protein |
| PA4469 | 145S | 0.104 | 1.41E-25 | 2.74E-14 | hypothetical protein |
| PA5438 | 274R | 0.08 | 6.33E-19 | 3.36E-14 | probable transcriptional regulator |
| PA3193 | 278L | 0.057 | 1.76E-11 | 1.52E-13 | glucokinase |
| PA2020 | 49Y | 0.115 | 1.06E-06 | 2.95E-13 | MexZ |
| PA3540 | 189E | 0.058 | 1.59E-15 | 3.72E-13 | GDP-mannose 6-dehydrogenase AlgD |
| PA4914 | 73A | 0.05 | 2.71E-14 | 7.30E-13 | transcriptional regulator, AmaR |
| PA3355 | 384D | 0.05 | 2.71E-14 | 3.48E-12 | hypothetical protein |
| PA3193 | 270T | 0.052 | 1.61E-13 | 3.79E-12 | glucokinase |

#### (1) PA5438

##### A. Identification

Mutations at amino acid loci 272-274 (SLR) of a probable transcriptional regulator PA5438 are significantly implicated in CF isolates (Table 2, highlighted in yellow). Fig. 3A demonstrates the amino acid residues of PA5438, and three residues underlined red presented outstandingly small lrt-pvalues amongst other amino acid residues. Amino acid sequence ‘SLR’ was deleted in 2419^th^ gene of the CF isolate 18A_661, a PA5438 homologue (Fig. 3B). Red lines in ‘id’ and rows ‘272^nd^’, ‘273^rd^’, ‘274^th^’ in Fig. 3C represent CF isolates, and deletions at the 272-274 loci, respectively. One exception of this is marked with an asterisk-arrow, where the 274^th^ amino acid residue R is replaced by C. SLR deletion in the PA5438 protein exhibits a highly positive correlation with CF isolates overall (Fig. 3C) and the PA5438 sequences except for the SLR sequence region were highly conserved when checked by multiple alignment of homologue sequences (data not shown). By performing NCBI protein domain search, these three amino acids were predicted to be a part of the end of the PRK11302 domain (data not shown) [41]. We constructed an in-frame deletion mutant, PA5438ΔSLR, to determine whether these three amino acids affect the function of the transcriptional regulator or not.

**Figure 3.**
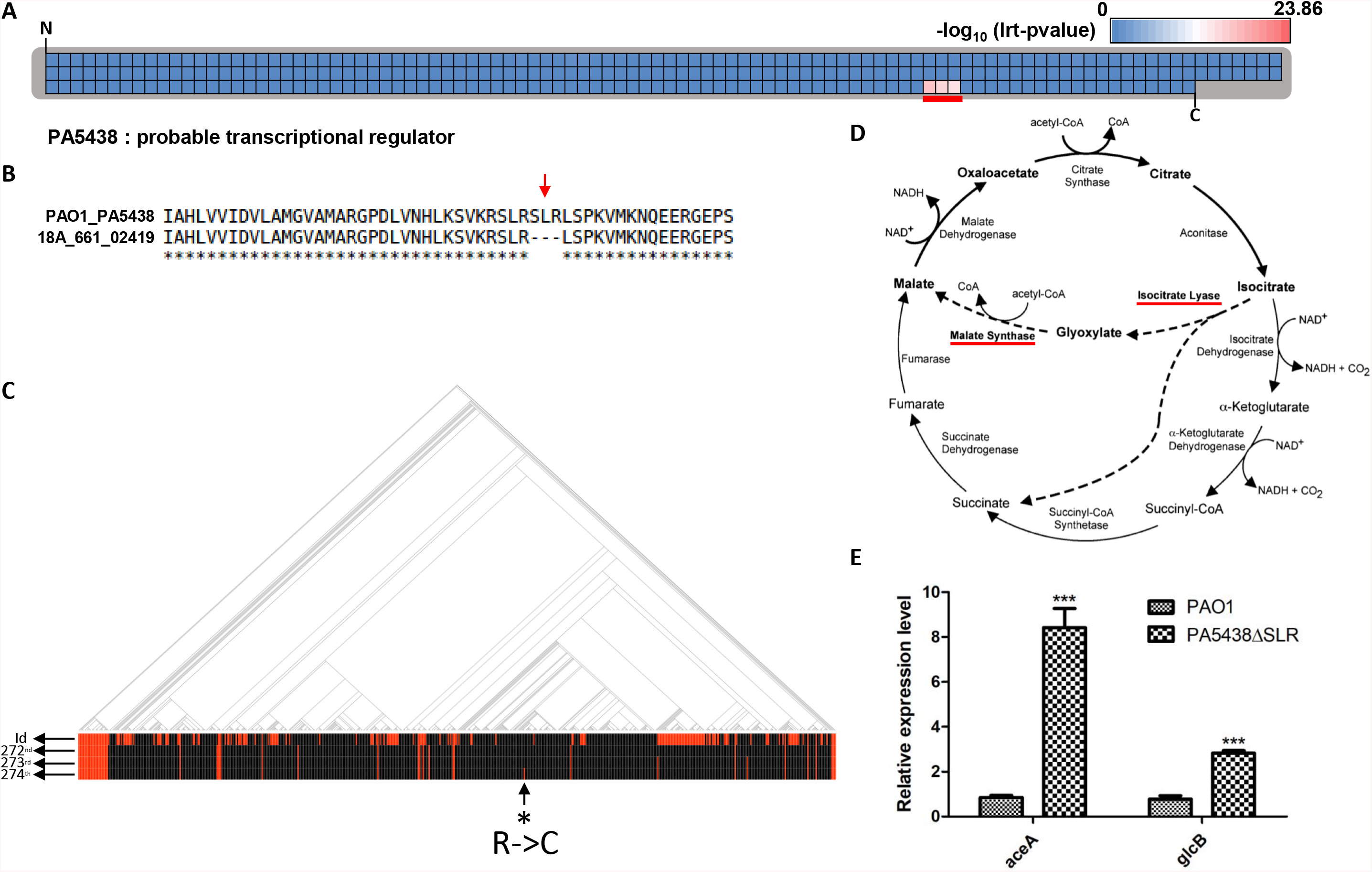
SLR deletion in PA5438 increases the expression level of *aceA* and *glcB*. **(A)** Total amino acid residues of PA5438 are presented and each amino acid locus is represented by individual squares. Colour of each square indicates the lrt-pvalue assigned to that amino acid residue based on Pyseer results and colours were assigned according to the index above the heatmap, which was generated based on the negative logs of the maximum and minimum lrt-pvalues from Pyseer result of amino acid locus. “N” and “C” mark the N- and C-terminus of PA5438, respectively. Residues presenting significantly low lrt-pvalues are marked with a red underline and were found at locus 272-274 of PA5438. **(B)** Comparison of the reference PA5438 to a homologue protein from a CF-isolated genome (18A_661_02419) is shown. Red arrow indicates deletion of SLR residues in PA5438 and corresponds to the red underlined locus observed in **(A)**. **(C)** Phylogenetic tree of 636 genomes and their associated disease status are shown. Red and black lines in the “id” row indicate genomes isolated from CF and non-CF patients, respectively. Red lines in the second, third and fourth rows (272nd, 273rd, 274th) each correspond to the absence of S, L, R residues, respectively, from the homologue. One exception to this is the red line in row 274th (asterisk-arrow), which was detected as R substituted by C. **(D)** AceA (isocitrate lyase) and GlcB (malate synthase) are enzymes involved in the glyoxylate shunt pathway (highlighted by red underlines). **(E)** RNAs of PAO1 and PA5438ΔSLR mutant were extracted at OD_600nm_ ~ 1.0 and relative expression levels of *aceA* and *glcB* were measured. \*\*\**p*<0.001

##### B. Experimental validation

PA5438 is a known suppressor which directly binds to the promoter region of *aceA* (isocitrate lyase) gene, and it has been shown to repress the expression of *glcB* (malate synthase) gene during growth in non-C2 carbon source [42]. These repressed genes encode enzymes involved in the glyoxylate shunt pathway (Fig. 3D) [43]. In order to determine whether the suppressive activity of PA5438 is lost in the PA5438ΔSLR mutant, gene expression levels of *aceA* and *glcB* were measured by Quantitative Real-Time PCR (qRT-PCR) using RNA extracted from bacterial cultures grown to OD_600nm_ ~1.0. qRT-PCR results show that the expression level increased 8-fold for *aceA* and 3-fold for *glcB*. Based on these findings, we expect that the deleted SLR sequence in PA5438 is a key region for determining the *aceA* and *glcB* expression levels (Fig. 3E).

Further investigation of the PA5438ΔSLR mutant was performed to identify phenotypes that may aid in the strain’s adaptation to the CF environment. First of all, we checked the phenotype of the mutant PA5438ΔSLR in LB media, as growth rates of diverse CF isolates in LB media closely approximated those of CF isolates grown in ASM (artificial sputum medium) and SCFM (synthetic CF sputum medium) [17]. Slower growth of the PA5438ΔSLR mutant compared to PAO1 was observed in LB, especially emphasised over the exponential phase (Fig. 4A). Since antibiotic tolerance can be caused by slowly-growing or non-dividing “persister” bacteria [44], we measured susceptibilities of PAO1 and PA5438ΔSLR mutant to popular antibiotics (tobramycin and ciprofloxacin) of different classes commonly used to treat *P. aeruginosa* infection in CF patients. In liquid culture with tobramycin or ciprofloxacin for 22 hours, PA5438ΔSLR mutant reached a significantly higher OD_600nm_ than PAO1 (Fig. 4B). When viable cells were counted after 22 hours, CFU of PA5438△SLR cultured in the presence of tobramycin showed a 10-fold increase compared to that of PAO1 (Fig. 4C), but there was no significant difference observed in CFUs of bacteria grown with shaking in the presence of ciprofloxacin (data not shown). A similar pattern of increased tolerance to tobramycin was observed in static bacterial culture, but no difference in susceptibility to ciprofloxacin was observed when measured by OD_600nm_ (Fig. S2A). However, when a higher concentration of ciprofloxacin was used (0.25 μg/ml), PAO1 growth was aborted while PA5438ΔSLR reached approximately OD_600nm_ ~ 0.15 after 18 hours of static culture (Fig. 4D). Therefore, we suspect that different mechanisms are involved in PA5438ΔSLR mutant’s increased tolerance to these two antibiotics.

**Figure 4.**
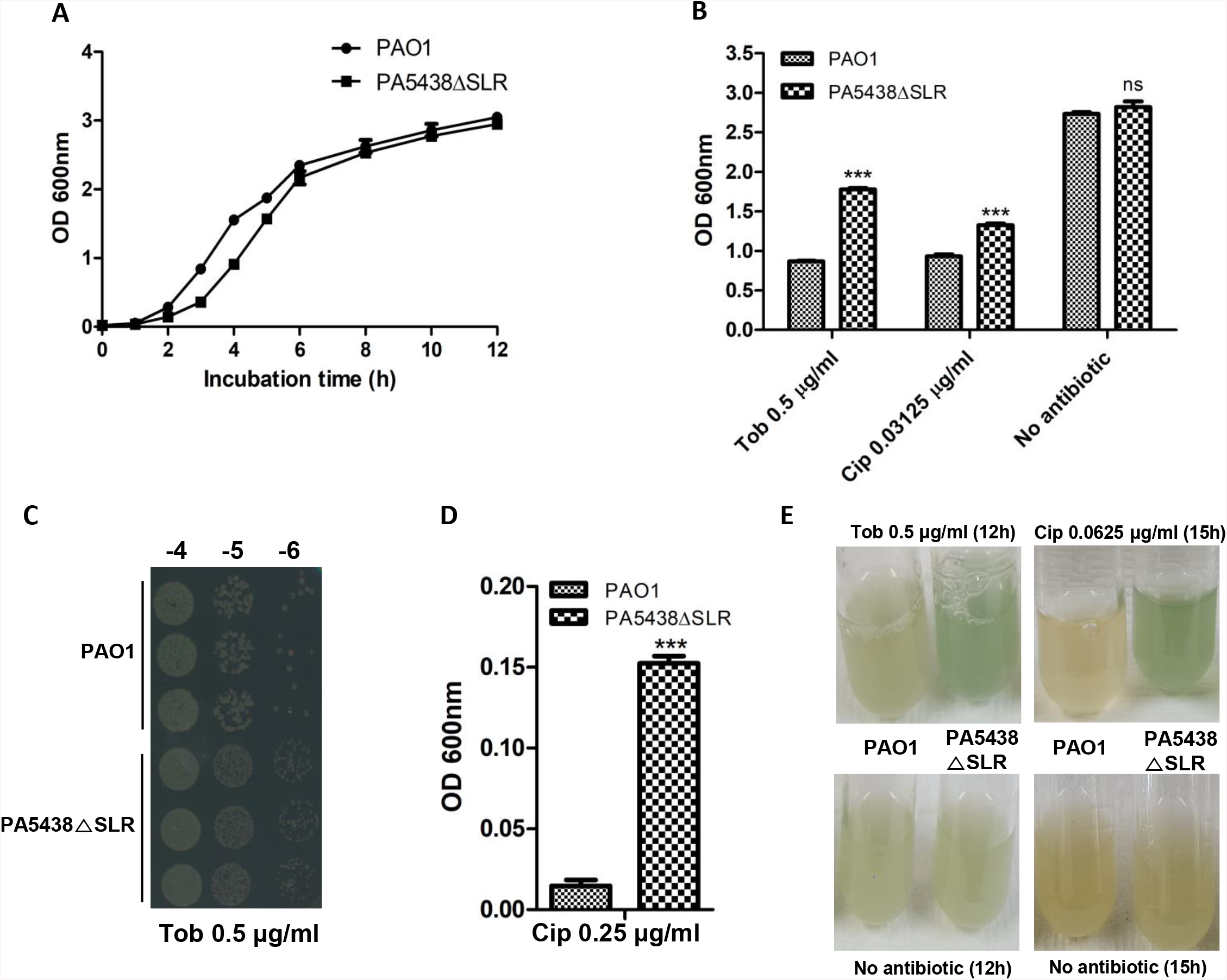
SLR deletion in PA5438 provide advantages under the antibiotic stress. **(A)** Growth curves of PAO1 and PA5438ΔSLR mutant in LB were observed over 12 hours. **(B)** Antibiotic susceptibility test against tobramycin (Tob) and ciprofloxacin (Cip) was performed. Initial CFU of PAO1 and mutant were adjusted to 5 × 105 CFU, and OD_600nm_ was measured after 22 hours of shaking incubation in LB supplemented with each antibiotic. Concentrations of antibiotics were 0.5 μg/ml and 0.03125 μg/ml, for tobramycin and ciprofloxacin, respectively. Growth with no antibiotic was measured as control. \*\*\**p*<0.001; ns: not significant **(C)** CFU of bacteria recovered from tobramycin susceptibility test was enumerated in triplicates on LB agar plates. The numbers above the pictures indicate the dilution factor used. **(D)** Antibiotic susceptibility tests with ciprofloxacin (Cip) were performed. Initial CFU of PAO1 and the mutant were adjusted to 5 × 105 CFU, and OD_600nm_ was measured after 18 hours of static incubation in LB supplemented with 0.25 μg/ml ciprofloxacin. \*\*\**p*<0.001 **(E)** Supernatant colours of PAO1 and the mutant cultures in LB after 12 hours with shaking incubation with 0.5 μg/ml tobramycin and no antibiotic are shown. In case of ciprofloxacin, 0.0625 μg/ml ciprofloxacin was used and bacteria were cultured for 15 hours with shaking incubation.

*P. aeruginosa* famously produces pyocyanin, an important virulence factor derived from phenazine and which forms as blue-green pigments. Pyocyanin induces ROS generation by transferring electrons to oxygen and increasing neutrophil apoptosis as a way of disrupting the host immune system [45]. As such, the supernatant colour of a *P. aeruginosa* culture can function as a proxy for the virulent nature of the bacterium. Interestingly, we observed the culture supernatant of PA5438ΔSLR under antibiotic stresses to be greener than that of the PAO1 culture supernatant (Fig. 4E). No such difference in the colours of culture supernatants was observed in the absence of antibiotics. Considering that the OD_600nm_ values of PAO1 and PA5438ΔSLR mutant cultures were not significantly different (Fig. S2B), we postulated that the PA5438ΔSLR mutant produces more pyocyanin and thus may be more virulent than PAO1 in the presence of tobramycin or ciprofloxacin.

##### C. Potential implications in CF airway infections

Altered fatty acid profile was found in the airways of CF patients compared with non-CF patients. For example, higher concentrations of palmitic acid and oleic acid were detected in the CF airway samples [46]. Fatty acid degradation upregulates the glyoxylate shunt (GS) pathway [47], and PA isolated from the CF lungs induced the expression of genes involved in fatty acid metabolism and GS pathway as determined by microarray [48]. Furthermore, mucin is a major energy source in the CF lung environment in addition to fatty acids, and the GS pathway is known to be important for mucin degradation and consumption [49]. Meta-transcriptomic analysis of several CF sputum samples in another study similarly found that *aceA* was upregulated and genes associated with glucose transporters and glycolysis were significantly downregulated [50]. Based on these previous studies, we assumed that activation of the GS pathway may provide an advantage to PA for proliferate in the CF lungs.

It seems likely that the previously observed upregulation of the GS pathway by microarray and meta-transcriptomic analysis is due to the presence of the CF isolates containing the abnormal PA5438 gene. In order to check whether the end products of the upregulated expressions of *aceA* and *glcB* are functional, we examined the sequences of AceA and GlcB from genomes that harbour ΔSLR. Both promoters and proteins of AceA and GlcB were highly conserved (Table S1, Table S2). Furthermore, we observed the growth of PA5438ΔSLR mutant in M9 media supplemented with glucose as sole carbon source was slower than PAO1 (Fig. S2C). Thus, we speculate that SLR deletion in PA5438 could be the cause of downregulated glucose metabolism. On the other hand, the mutant grew slightly better in M9 with acetate as the sole carbon source, a condition wherein the GS pathway is essential for bacterial growth [51] (Fig. S2D).

Based on our experimental results, we speculate that the appearance of this mutant may present problems of increased tolerance to tobramycin and ciprofloxacin, and increased virulence in CF-associated PA infection. Further studies will be conducted to reveal the mechanisms of increased tolerances to tobramycin and ciprofloxacin, and the accompanied increase in virulence.

#### (1) YecS (PA0313)

##### A. Identification

Top 20 amino acid residues whose corresponding 31mers do not align to the PAO1 genome are described in Table 3. A major limitation of aligning the 31mers to the PAO1 genome as reference, is that, 31mers that fail to align to the PAO1 genome as result of insertion mutations are overlooked. Hence, additional analysis was performed to identify insertion mutations significantly implicated in either CF or non-CF isolates. In order to determine the reference genes for such insertion mutations, 31mers that failed to align to the PAO1 genome were *de novo* assembled, and the contigs that formed were aligned to 635 genomes (PAO1 was excluded). For instance, the contig constructed from 31mers that included 9 extra nucleotides encoding the additional SLI sequence, highlighted in yellow (Table 3), aligned to the 4951^st^ protein of AU17965_3981 (a YecS homologue). Hence, AU17965_3981_04951 was selected as the reference protein.

**Table 3.**
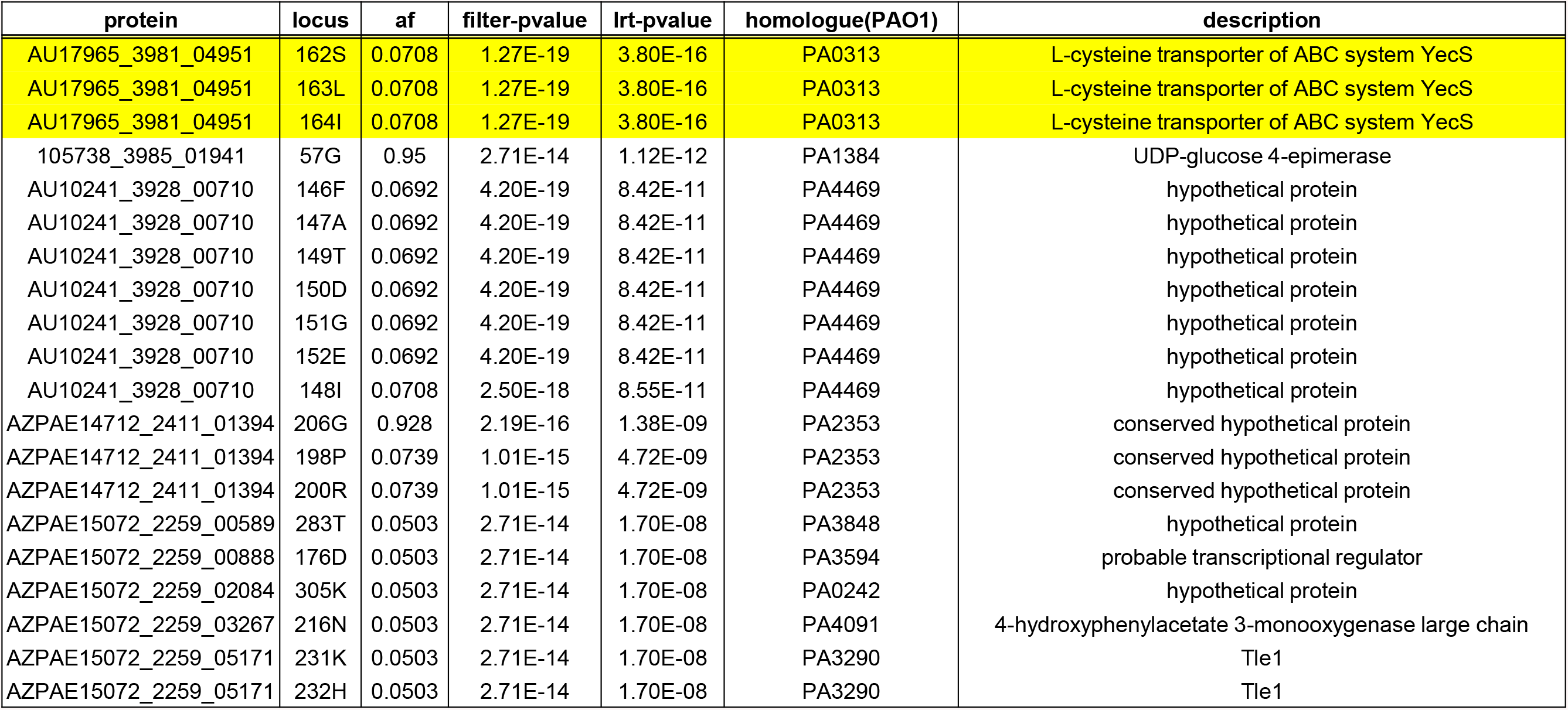
Top 20 amino acid residues of the small variant group whose 31mers did not align to the PAO1 genome. Top 20 amino acid residues according to Pyseer results whose 31mers did not align to the PAO1 genome are shown. Reference protein from genomes other than PAO1 and the PAO1 homologue of this reference protein are presented in columns “protein” and “homologue”. Column “locus” contains locus of the mutation and amino acid of the reference protein where the mutation is detected. Multiple types of mutation at each locus of the reference protein may be present and detailed information is provided in supplementary data 3. The reference protein name in the “protein” column is composed of the genome id, connected by the latter underscore sign, to gene number within that genome. All sequences of the reference proteins in Table 3 are provided in supplementary data 4.

| protein | locus | af | filter-pvalue | lrt-pvalue | homologue(PAO1) | description |
| --- | --- | --- | --- | --- | --- | --- |
| AU17965_3981_04951 | 162S | 0.0708 | 1.27E-19 | 3.80E-16 | PA0313 | L-cysteine transporter of ABC system YecS |
| AU17965_3981_04951 | 163L | 0.0708 | 1.27E-19 | 3.80E-16 | PA0313 | L-cysteine transporter of ABC system YecS |
| AU17965_3981_04951 | 164I | 0.0708 | 1.27E-19 | 3.80E-16 | PA0313 | L-cysteine transporter of ABC system YecS |
| 105738_3985_01941 | 57G | 0.95 | 2.71E-14 | 1.12E-12 | PA1384 | UDP-glucose 4-epimerase |
| AU10241_3928_00710 | 146F | 0.0692 | 4.20E-19 | 8.42E-11 | PA4469 | hypothetical protein |
| AU10241_3928_00710 | 147A | 0.0692 | 4.20E-19 | 8.42E-11 | PA4469 | hypothetical protein |
| AU10241_3928_00710 | 149T | 0.0692 | 4.20E-19 | 8.42E-11 | PA4469 | hypothetical protein |
| AU10241_3928_00710 | 150D | 0.0692 | 4.20E-19 | 8.42E-11 | PA4469 | hypothetical protein |
| AU10241_3928_00710 | 151G | 0.0692 | 4.20E-19 | 8.42E-11 | PA4469 | hypothetical protein |
| AU10241_3928_00710 | 152E | 0.0692 | 4.20E-19 | 8.42E-11 | PA4469 | hypothetical protein |
| AU10241_3928_00710 | 148I | 0.0708 | 2.50E-18 | 8.55E-11 | PA4469 | hypothetical protein |
| AZPAE14712_2411_01394 | 206G | 0.928 | 2.19E-16 | 1.38E-09 | PA2353 | conserved hypothetical protein |
| AZPAE14712_2411_01394 | 198P | 0.0739 | 1.01E-15 | 4.72E-09 | PA2353 | conserved hypothetical protein |
| AZPAE14712_2411_01394 | 200R | 0.0739 | 1.01E-15 | 4.72E-09 | PA2353 | conserved hypothetical protein |
| AZPAE15072_2259_00589 | 283T | 0.0503 | 2.71E-14 | 1.70E-08 | PA3848 | hypothetical protein |
| AZPAE15072_2259_00888 | 176D | 0.0503 | 2.71E-14 | 1.70E-08 | PA3594 | probable transcriptional regulator |
| AZPAE15072_2259_02084 | 305K | 0.0503 | 2.71E-14 | 1.70E-08 | PA0242 | hypothetical protein |
| AZPAE15072_2259_03267 | 216N | 0.0503 | 2.71E-14 | 1.70E-08 | PA4091 | 4-hydroxyphenylacetate 3-monooxygenase large chain |
| AZPAE15072_2259_05171 | 231K | 0.0503 | 2.71E-14 | 1.70E-08 | PA3290 | Tle1 |
| AZPAE15072_2259_05171 | 232H | 0.0503 | 2.71E-14 | 1.70E-08 | PA3290 | Tle1 |

Lrt-pvalues of the individual amino acid residues of AU17965_3981_04951 protein are depicted in Fig. 5A. Three amino acid residues at loci 162-164, indicated by a red underline, had significant lrt-pvalues compared to other amino acid residues. At these loci, the YecS homologue had insertions of three amino acid residues, SLI (Fig. 5B). Moreover, there was an additional copy of SLI in the AU17965_3981_04951 protein, comprising a total of three stretches of ‘SLI’, in contrast to a total of two stretches in the PAO1 YecS (Fig. S3). No other differences in the amino acid sequences were detected between the two proteins. Strong positive correlation between CF isolates (represented by red coloured “id” lines) and insertion of ‘SLI’ (indicated by red coloured “162^nd^”,”163^rd^”, “164^th^” lines) was detected across the genomes illustrated in the phylogenetic tree (Fig. 5C). Furthermore, the YecS homologue sequences of these SLI insertion-carrying CF isolates were highly conserved, except for the additional ‘SLI’, in comparison to the PAO1 YecS (data not shown). AU17965_3981_04951 protein is a cytoplasmic membrane transporter, and the additional ‘SLI’ sequence was predicted to span both cytoplasmic and transmembrane regions (Fig. S4A, Fig. S4B), by Phobius [52].

**Figure 5.**
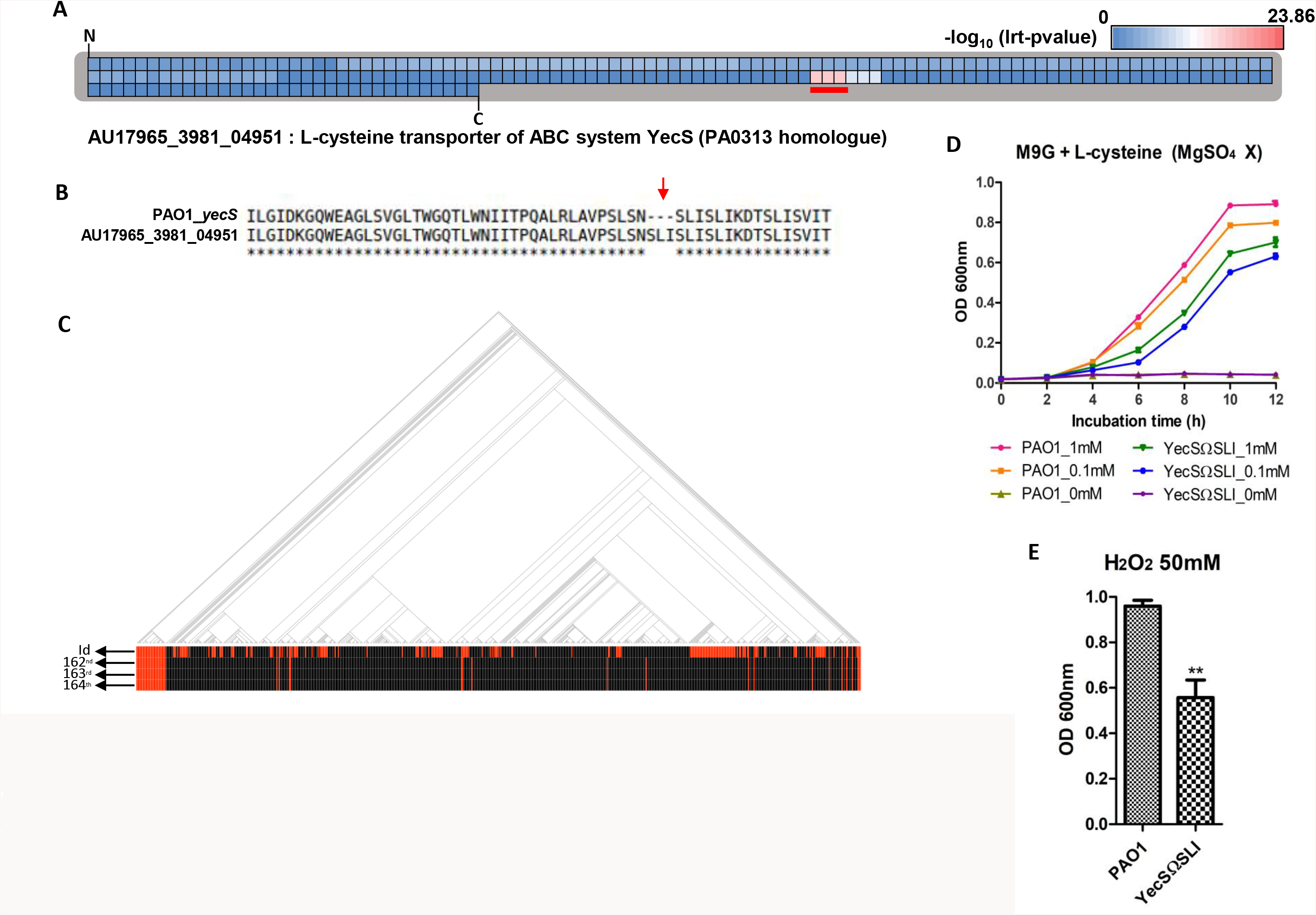
SLI insertion in YecS reduces the L-cysteine utilization activity and resistance to hydrogen peroxide. **(A)** Total amino acid residues of CF-derived AU17965_3981_04951 protein are presented and each amino acid locus is represented by individual squares. Colour of each square indicates the lrt-pvalue assigned to that amino acid residue based on Pyseer results and colours were assigned according to the index above the heatmap. The colour index was generated based on the negative logs of the maximum and minimum lrt-pvalues from Pyseer result of the amino acid locus. “N” and “C” mark the N- and C-terminus of AU17965_3981_04951, respectively. Residues presenting especially low lrt-pvalues are marked with a red underline and were found at loci 162-164 of the AU17965_3981_04951. **(B)** Comparison of the reference AU17965_3981_04951 to its PAO1 homologue YecS (PA0313) is shown. Red arrow indicates insertion of SLI residues in AU17965_3981_04951 and corresponds to the red underlined locus observed in **(A)**. **(C)** Phylogenetic tree of 636 genomes and their associated disease status are shown. Red and black lines in the “id” row indicate genomes isolated from CF and non-CF patients, respectively. Red lines in the second, third and fourth rows indicate the presence of SLI insertion in that homologue. **(D)** Growth curves of PAO1 and YecSΩSLI mutant cultured in M9 minimal media with no MgSO_4_, supplemented with glucose and L-cysteine (1 mM, 0.1 mM, 0 mM) as the sole sulphur source were measured over 12 hours. **(E)** OD_600nm_ of PAO1 and YecSΩSLI mutant in LB with 50 mM hydrogen peroxide were measured after 6 hours of shaking incubation. \*\**p*<0.01

##### B. Experimental validation

In order to find out whether the ‘SLI’ insertion results in any phenotypic change, we constructed an in-frame mutant with the additional ‘SLI’ sequence inserted into the corresponding locus of PAO1 YecS. No difference in growth was observed between the wild type PAO1 and the YecSΩSLI mutant when both strains were grown in LB (Fig. S4C). However, growth in M9 minimal media supplemented with glucose and L-cysteine as the sole sulphur source showed significant difference between PAO1 and the mutant, whereas growth in M9 media with an additional sulphur source present (MgSO_4_) showed no difference (Fig. 5D, Fig. S4D).

In *E. coli*, two L-cystine transporters exist; ATP-binding cassette (ABC) importer FliY-YecSC and symporter YdjN [53]. In a study by Ohtsu et al. (2015), both YecS and YdjN single deletion mutants exhibited the same growth rate as wild type in M9 minimal media containing glucose and L-cystine as the sole sulphur source. In contrast, a double deletion mutant showed no growth in the same media condition [53]. These findings imply that the presence of either of the two transporters is sufficient to meet the L-cystine requirement for normal growth when L-cystine is used as the sole sulphur source. In our study, we postulated that L-cysteine is an appropriate alternative to L-cystine for our purposes, as L-cysteine reacts with periplasmic hydrogen peroxide and gets converted into L-cystine [54]. Unlike *E. coli*, *P. aeruginosa* does not seem to encode a YdjN homologue based on our web-based Uniprot search, since no PA genomes were detected to contain proteins functionally annotated as YdjN [55]. Therefore, based on our Uniprot search and our experimental result (Fig. 5D), we hypothesise that PAO1 possesses only the ABC importer system for L-cysteine uptake. Moreover, the decreased growth of the SLI insertion mutant is attributed to a decreased activity of the ABC transporter rather than a complete loss of function. Comparison of growth between a YecS clean deletion mutant and our SLI insertion mutant would be helpful to better elucidate the assortment of L-cysteine transporters present in *P. aeruginosa*.

Previous studies by Ohtsu et al., also found a FliY clean deletion and a double deletion mutant of YecS and YdjN exhibited increased hydrogen peroxide sensitivity as the L-cysteine/L-cystine shuttle system is disrupted in these *E. coli* mutants [53, 54]. For this reason, we hypothesised that a malfunctional L-cysteine transporter may influence hydrogen peroxide resistance of *P. aeruginosa*. To test this idea, hydrogen peroxide resistance test was performed by challenging the YecSΩSLI mutant with 50 mM hydrogen peroxide in LB. Indeed, the mutant exhibited decreased growth compared to PAO1 (Fig. 5E).

##### C. Potential implications in CF airway infections

In addition to the SLI insertion mutation that we focused on, other patterns of mutation were observed across the CF isolate genomes. One such pattern is the deletion of a ‘SLI’ sequence at the locus indicated by the asterisk in **a** of Fig. S3, and this results in CF isolates encoding just one ‘SLI’ sequence in their YecS homologues (Fig. S3). Another pattern of mutation observed by multiple alignment of YecS homologues is deletion of long stretches of amino acid residues (Fig. S3). Since the SLI insertion mutant exhibited a decreased L-cysteine transporter activity (Fig. 5D), we anticipate that such large deletions in the YecS protein would incur in a complete loss of function. It is possible that hydrogen peroxide sensitivity of CF isolates with large alterations in YecS is increased to a greater degree than that of the YecSΩSLI mutant.

Thick dehydrated mucus layer caused by mutated CFTR is known to establish a microaerobic or even anaerobic environment in the CF airway [56]. O_2_ concentration decreases steeply from the mucus layer to the respiratory epithelium, and obligate anaerobic bacteria have been detected in the anaerobic CF lung [56, 57]. Within this anaerobic environment, PA implements survival strategies like denitrification and fermentative pathway to produce energy, as an alternative to respiration which generates reactive oxygen species (ROS). Thus, we speculate that the susceptibility to H_2_O_2_ of YecSΩSLI would not confer a significant disadvantage in dwelling in the CF airway, and further studies are required to understand the implications of this mutation under anaerobic conditions.

### Small-scale alterations in intergenic regions: *phuS* and *phuR*

Comparison of genomes using k-mer analysis is advantageous in that it enables the investigation of intergenic regions. Of all contigs assembled from significantly different 31mers between CF and non-CF groups, 29 contigs aligned to the non-coding regions of the PAO1 genome. The regions that most of these 29 contigs bind to are either contained within the hs region, or within rDNA sequences, as illustrated in Fig. 1A. Since we suspect that mutations located in the hs region to cause gene deletions, and rDNA sequence regions were excluded from analysis due to difficulties in interpretation, we selected regions other than these loci. Amongst several such intergenic regions, we found one intergenic region between *phuR* and *phuS* operons which are involved in the pseudomonas heme utilization (phu) systems (Fig. 6A). The function of such system is the acquirement of iron from the heme group of hemoglobin [58]. In order to evaluate the potential role of these mutations in CF adaptation, we performed Pyseer analysis using the nucleotide sequences of this intergenic region from PAO1 and other isolates as input. Top 5 mutations in the increasing order of lrt-pvalues are listed in Fig. 6B. The top hit with the smallest lrt-pvalue in this list is the transition of the 117^th^ residue on the forward strand from cytosine (C) to thymine (T) (Fig. 6B). Two mutations (C117T, C122T) are included in the *phuR* promoter region (from −35 region to transcriptional initiation site (+1)) (Fig. 6C).

**Figure 6.**
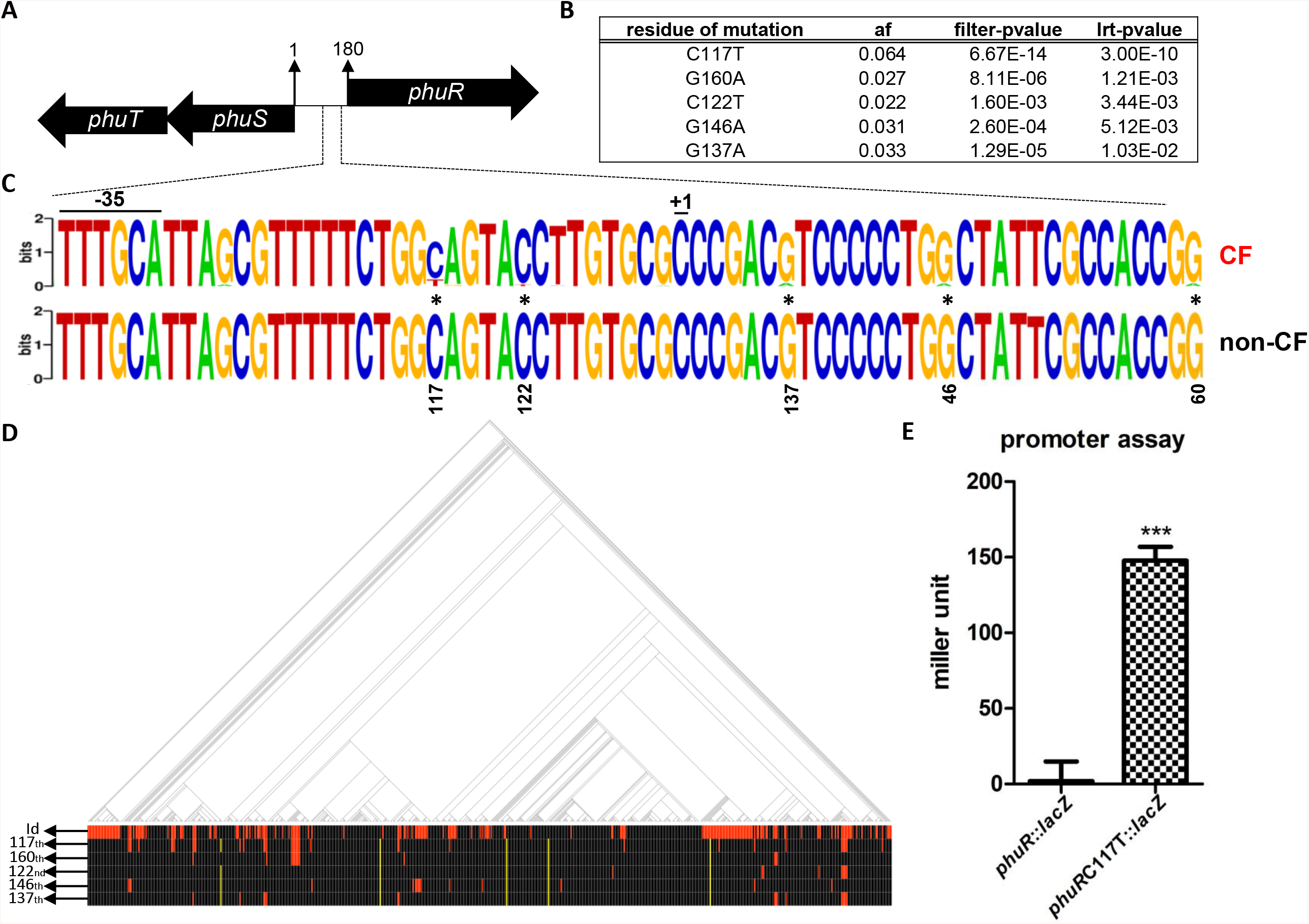
One SNP in *phuR* promoter region increases expression level. **(A)** Region encompassing the intergenic region of *phuR* and *phuS* is shown, and the length of the intergenic region is 180 bp. **(B)** Top 5 SNPs detected by Pyseer within this intergenic region are listed. The number in the “residue of mutation” column indicates the forward strand-based locus of mutation. The nucleotide in front of the locus number indicates the reference base in the intergenic region, whereas the nucleotide following the locus number indicates the changed base. Information contained in the following three columns are similar to those described in Fig. 2B. **(C)** Promoter region of *phuR* (region from −35 to +1) based on PRODORIC database is shown, and SNPs listed in **(B)** are marked by asterisks. The height of a stack reflects the degree of sequence conservation at that position, while the height of symbols within a stack indicates the relative frequency of each nucleotide at that position. The Upper sequence is derived from the intergenic regions of CF genomes and sequence below is drawn based on intergenic regions of non-CF genomes. **(D)** Red lines in the bottom four rows indicate mutations at each locus, shown in **(B)**, and yellow lines indicate that this intergenic region is missing from the genome. **(E)** Culture of *phuR*::*lacZ* and *phuRC117T*::*lacZ* grown to OD_600nm_ ~0.25 were used in β-galactosidase assay, to compare *phuR* promoter activity. \*\*\**p*<0.001

Consistent with our findings, frequency of mutations within this intergenic region in a CF isolate was detected to be increased significantly compared to the expected mutation rate by chance, and the mutated intergenic region increased the *phuR* promoter activity [59]. Yellow lines in Fig. 6D indicate the deletion of the intergenic region, and thus the promoter activity is probably absent in these cases. When such cases of promoter deletion are excluded from consideration, a stronger correlation between the mutations of this region and CF isolates is expected, and thus lrt-pvalues are expected to decrease. To examine whether C117T point mutation affects the expression of *phuR*, we performed a promoter activity assay by measuring β-galactosidase activity. Promoters with this mutation exhibited increased activity (Fig. 6E). Therefore, we anticipate that the increased *phuR* promoter activity to ultimately result in increased phu system activity. We speculate that C117T mutation is an important strategy for iron uptake and thus survival in the CF environment.

## Conclusions

In this work, we compared a large set of genomes from clinical PA isolates (CF vs. non-CF) and identified genetic mutations that specifically occurred in either CF or non-CF isolates. These findings were made possible only because we performed GWAS based on 31mer counting. We also wondered if the DNA mutations that we specified would indeed impact bacterial growth-associated phenotypes. Importantly, PAO1-derived variants, PA5438ΔSLR and YecSΩSLI, exhibited distinct phenotypes in our *in vitro* assays. Moreover, a single nucleotide replacement in the promoter region of *phuR* gene caused a robust increase in the gene transcription. Together, these results suggest that PA might take advantage of small-scale mutations when establishing chronic infections in the CF airway.

Mutations in *lasR* gene encoding a quorum sensing (QS) regulator were reported to be frequently seen in CF isolates [7, 8]. In our study, however, mutations in *lasR* or other QS-related genes were identified not to be statistically significant between CF and non-CF isolates. Given that the *mucA* mutation was clearly represented in our analysis, which demonstrates the integrity of our bioinformatic approach, this information suggests that *lasR* mutation might be a common feature in PA either causing chronic CF infection or other types of acute infections.

PA infection has been a very important medical issue. In order to control its widespread and recalcitrant infection, it is crucial to understand how PA evolves its genome to adapt to specific host environments. Our results will stimulate further investigations to better understand which genes and promoter regions in PA genomes are specifically targeted for alterations in chronic CF infections.

## Experimental Procedures

### Bioinformatic analysis

2,187 *P. aeruginosa* genomes were downloaded from the Pseudomonas Genome Database [25], and 1,001 genomes with information of host disease were selected. Phylogenetic tree was drawn by RapidNJ [26] and visualized by Microreact [60]. Based on the tree generated, the selection of genomes was trimmed down by Treemmer [27] to contain 636 isolates while 99.8 % of original diversity was maintained (supplementary data 1). Prediction of protein-coding genes of 636 genomes were performed using Prokka [32]. K-mers 31 base pairs long (31mer) were counted in 636 individual genomes by Fsm-lite [30], and a similarity matrix was constructed by Snp-sim [61] using a core alignment of 636 genomes generated by Snippy [62]. Pyseer [29], which employs a mixed model (FaST-LMM), was run using the counted k-mers and the similarity matrix as input, and lrt-pvalues were assigned to each 31mer. In order to sort 31mers significantly different between CF and non-CF groups, 31mers with bad-chi values or lrt-pvalues above 2.8e-08 (cutoff value in this analysis) were removed. As a result, 41,685 31mers were detected to have significant associations with either disease statuses (CF or non-CF). Distribution of these 31mers aligned to the PAO1 whole genome was visualized with Phandango [31]. Subsequently, *de novo* assembly of these 31mers was completed by Trinity [33]. 494 contigs were constructed, and these contigs aligned to 29 intergenic regions, 6 untranslated regions and 109 translated genes by Blastn. In case of 109 contigs binding to protein coding regions, locus information was used to derive amino acid sequences from the nucleotide sequences. Individual contigs binding to both protein-coding and intergenic regions were blasted against 635 genomes (PAO1 genome was excluded) [34]. Top hit from each pairwise alignment with e-value smaller than 0.01 was chosen as homologue of the candidate sequence in each isolate. If no significant hit was retrieved, we assumed that homologue corresponding to the candidate region is deleted in the isolate genome. Multiple alignment of the candidate sequence and its homologues was executed with Mafft [63] to check the nucleotide (in case of intergenic region candidates) or amino acid sequence (in case of protein-coding genes) at each locus. If the residue of a homologue was identical to that of the reference intergenic region or protein, it was assigned a value of ‘1’ at that locus, and if not, a value of ‘0’ was assigned. In this fashion, a locus matrix was obtained.

Subsequently, protein-coding homologues were clustered under several conditions of similarity and coverage using Blastclust [34]. Again, homologues that got clustered with the reference protein was assigned ‘1’ and those that did not was assigned ‘0’. We then used the clustering matrix and the aforementioned locus matrix as input for Pyseer. Loci and clusters with either bad-chi values or lrt-pvalues larger than 4.01E-06 and 3.22E-05, respectively, were removed. In this manner, we were able to determine meaningful variants (Fig. 1B) and loci. Additionally, based on the number of meaningful loci implicated in a candidate protein, the mutation was classified either as a small or large variant; if % of variable loci is greater than 20% of the protein length, the protein was categorised as a large variant, and if not, a small variant. Finally, k-mers that failed to align to the PAO1 genome were also similarly analysed. However, the PAO1 reference gene in these cases was replaced with the reference gene detected in alternative genomes. As such, investigation of insertion variants was made possible. For KEGG functional annotation and pathway analysis, BlastKoala [35] was performed with the large variants. Sequence comparison was performed by Clustalw [64] and multiple alignment was visualised with Jalview [65].

### Bacterial strains and growth conditions

All bacterial strains and plasmids used in this study are shown in Table S3. *Pseudomonas aeruginosa* PAO1 was used as a reference strain and all in-frame mutants including SLI insertion in YecS protein, SLR deletion in PA5438 protein were constructed from the PAO1 strain. Bacterial cultures were grown in Luria-Bertani (LB) medium (1% [w/v] tryptone, 0.5% [w/v] yeast extract, and 1% [w/v] sodium chloride) at 37 °C. All bacterial single colonies on LB plates were picked and inoculated in LB broth for precultures and grown overnight. Precultures were diluted 100-fold in fresh LB broth for subculture and incubated at 37 °C with shaking at 230 rpm. The incubation time was dependent on the experimental procedures. *E. coli* used in the cloning process also used LB broth or LB broth supplemented with 50 μg/ml of gentamicin (Sigma-Aldrich, USA) and 30 μg/ml of ampicillin (Sigma-Aldrich, USA). For screening the single crossover recombinants, LB agar plates with 50 μg/ml of gentamicin and 20 μg/ml irgasan (Sigma-Aldrich, USA) were used, and LB agar plates without NaCl but with 8% sucrose were used to select insertion and deletion mutants. In antibiotic susceptibility test under shaking bacterial culture, 0.5 μg/ml tobramycin (Sigma-Aldrich, USA), 0.0625 μg/ml and 0.03125 μg/ml ciprofloxacin (Duchefa, Netherland) were supplemented in LB, and under static bacterial culture, 1 μg/ml tobramycin, and 0.25 μg/ml and 0.0625 μg/ml ciprofloxacin were supplemented in LB.

### In-frame mutant construction

In-frame insertion (SLI insertion in of YecS protein) and deletion (SLR deletion of PA5438) were performed to construct mutants with amino acid level changes. In case of the in-frame deletion, 5’ and 3’ flanking regions in both direction of SLR were designed to overlap. However, for in-frame insertion, nucleotide sequences corresponding to amino acid sequence ‘SLI’ were inserted into the middle of the 5’ flanking region. Overlap was constructed by using each 5’ primer of 5’ flanking region and 3’ primer of 3’ flanking region. For both deletion and insertion mutations, overlapping product was inserted into modified pCVD442, a suicide vector, containing the gentamicin and ampicillin resistance markers. PAO1, grown on LB agar, was conjugated with *E. coli* SM10 λpir, harbouring pCVD442 with the overlap product inserted, grown on LB agar with 50 μg/ml gentamicin and 30 μg/ml ampicillin. Conjugates were spread onto LB agar with 50 μg/ml gentamicin and 20 μg/ml irgasan to select single crossover recombinants. This single crossover recombinant was incubated on LB agar without NaCl but containing 8% sucrose for selection of the desired mutant. Sequence verification was performed by PCR. Primers used in constructing these mutants are listed in Table S4.

### Promoter assay

In order to perform *phuR* gene promoter assay, intergenic regions (180 bp) from the PAO1 genome and AU2342_3932 [66] with only the 117^th^ locus changed from cytosine to thymine were amplified with primers listed in Table S4. These PCR-products were each cloned into the upstream region of the β - galactosidase gene of puc18-mini-Tn7t-Gm-LacZ [67] for chromosomal insertion. The constructed plasmid was transformed into *E. coli* DH5α λpir. After verification by DNA sequencing, this plasmid with the helper plasmid pTNS2 that encodes TnsABC+D genes, which allow Tn7 transposition, were electroporated into PAO1. Empty puc18-mini-Tn7t-Gm-LacZ plasmid with no insert was used as control to measure the baseline expression of *lacZ*. The potential clones were selected on LB agar with 50 μg/ml gentamicin and sequence verification was performed to select the final candidates whose transposon was inserted properly into region following *glmS* gene. β - galactosidase activities of three clones (con::*lacZ*, *phuR*::*lacZ* and *phuR*C117T::*lacZ*), summarised in Table S3, were measured at exponential phase (OD_600nm_ ~ 0.25) grown in LB.

### Growth curves

PAO1, YecS and PA5438 mutants were precultured overnight in LB broth, diluted 100-fold in fresh LB and incubated at 37 °C with shaking at 230 rpm. Growth in LB was observed over a period of 12 hours and OD_600nm_ was measured. To examine the capacity of the L-cysteine transporter, overnight pre-cultures of PAO1 and the YecS mutant were washed in phosphate-buffered saline (PBS) and diluted 100-fold in 1X M9 minimal media supplemented with 22.2 mM glucose, and varying concentrations of L-cysteine (1 mM, 0.1 mM, 0 mM) as sole sulphur source (without MgSO_4_). OD_600nm_ was measured over 12 hours. Growth was also measured over 10 hours under identical conditions, except for the inclusion of 2 mM MgSO_4_ in 1X M9 minimal media. For PA5438 mutant culture, 1X M9 media supplemented with carbon source (22.2 mM glucose or 66.6 mM sodium acetate) was used to monitor growth over 14 hours.

### Hydrogen peroxide resistance test

Precultures of PAO1 and the YecS mutant for the hydrogen peroxide resistance test were prepared in the same manner as described above. Precultures were subsequently diluted 100-fold in fresh LB containing 50 mM hydrogen peroxide (Sigma-Aldrich, USA). After incubation at 37 °C with shaking at 230 rpm for 6 hours, OD_600nm_ was measured.

### Antibiotic susceptibility test and state of culture supernatant under antibiotic stress

For antibiotic susceptibility test, ciprofloxacin and tobramycin were used. PAO1 and PA5438 mutant were precultured overnight in LB broth. Bacterial preculture was diluted 100-fold in fresh LB and this was incubated at 37 °C with shaking at 230 rpm for 3 and 4 hours, for PAO1 and the PA5438 mutant respectively, to adjust for the difference in growth rates.

These bacterial cultures were adjusted to 5 × 10^5^ colony-forming units (CFU) and incubated overnight in LB supplemented with ciprofloxacin and tobramycin. For shaken cultures, antibiotic concentrations used were: 0.5 μg/ml tobramycin, 0.03125 μg/ml ciprofloxacin. For static cultures, antibiotic concentrations used were: 1 μg/ml tobramycin, 0.0625 μg/ml and 0.25 μg/ml ciprofloxacin. After 22 hours of shaking incubation, the numbers of viable cells were counted by spotting 10-fold serial dilutions of the bacterial culture on LB agar plates, and incubating overnight at 37 °C. Colours of supernatants collected from shaken cultures under several antibiotic conditions, 0.0625 μg/ml ciprofloxacin and 0.5 μg/ml tobramycin, or no antibiotic stress, were recorded after 12 and 15 hours. OD_600nm_ was also measured.

### Reverse Transcription and Quantitative Real Time PCR

PAO1 and PA5438 mutant were precultured and subcultured in LB. After incubating the subcultures to OD_600nm_ ~ 1.0, RNeasy Mini kit involving on-column DNase1 digestion (Qiagen, Netherland) was used following the manufacturer’s protocol to extract RNA. 1 μg of RNA was reverse-transcribed to synthesize complementary DNA by using reverse transcriptase (Takara Bio, Japan) and random hexamer primers. To check for DNA contamination, the same process of cDNA synthesis was performed in the absence of reverse transcriptase. SYBR-green-based qPCR was performed using ABI 48-well StepOne™ real-time system, and the primers used are listed in Table S4. Annealing was done at 66 °C and CT values were normalized by 16S rRNA CT values.

### Statistical Analysis

Data shown are expressed as means ± standard deviation. Unpaired Student’s t-test (one-tailed, unequal variance) was performed to analyse the differences between experimental groups. P-values smaller than 0.05 were considered statistically significant. All experiments were repeated for reproducibility.

## Acknowledgements

This work was supported by grants from the National Research Foundation (NRF) of Korea, which is funded by the Korean Government (2017M3A9F3041233 and 2019R1A6A1A03032869). This research was also supported by a grant of the Korea Health Technology R&D Project through the Korea Health Industry Development Institute (KHIDI), funded by the Ministry of Health & Welfare, Republic of Korea (HI14C1324).

## Author Contributions

W.H. and S.S.Y. conceptualized and designed the experiments. W.H. performed experiments and analyzed experimental results. W.H., J.H.Y. and S.S.Y. drafted the manuscript. J.H.Y., K.B.M., and K.L. provided valuable comments. All authors read and approved the final manuscript.

---

**Figure S1.**
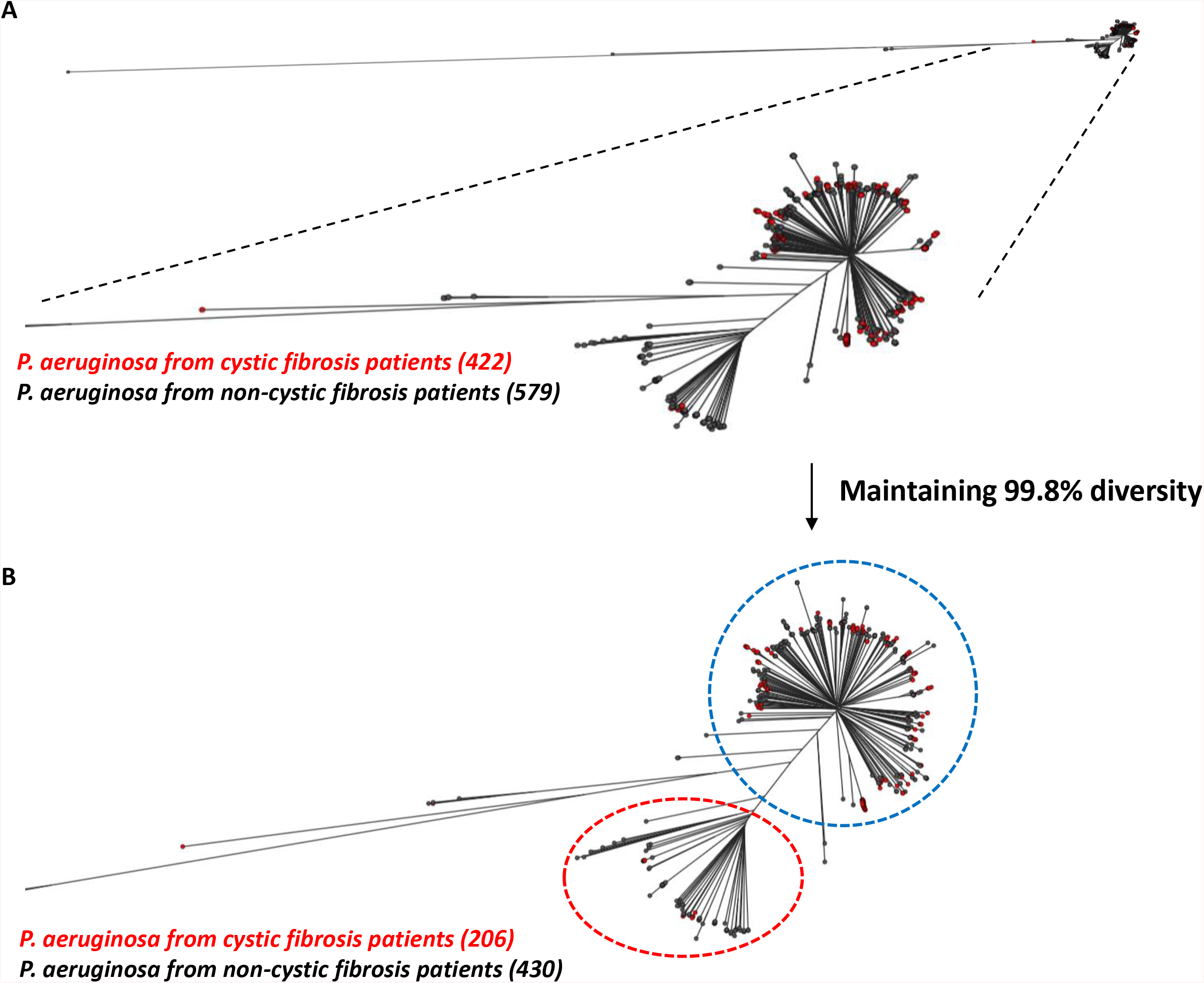
Phylogenetic tree constructed with CF and non-CF genomes. Upper phylogenetic tree was constructed with 1,001 genomes containing host disease information, and the tree below was drawn using 636 genomes and maintaining 99.8% diversity of the upper tree. Black and red leaves each indicate non-CF and CF isolates, and the numbers of CF and non-CF genomes for constructing each phylogenetic tree are placed inside brackets. Most of the CF isolates are located in the main group (blue dotted line) which includes the majority of genomes, and the smaller group is referred to as the sub-group (red dotted line).

**Figure S2.**
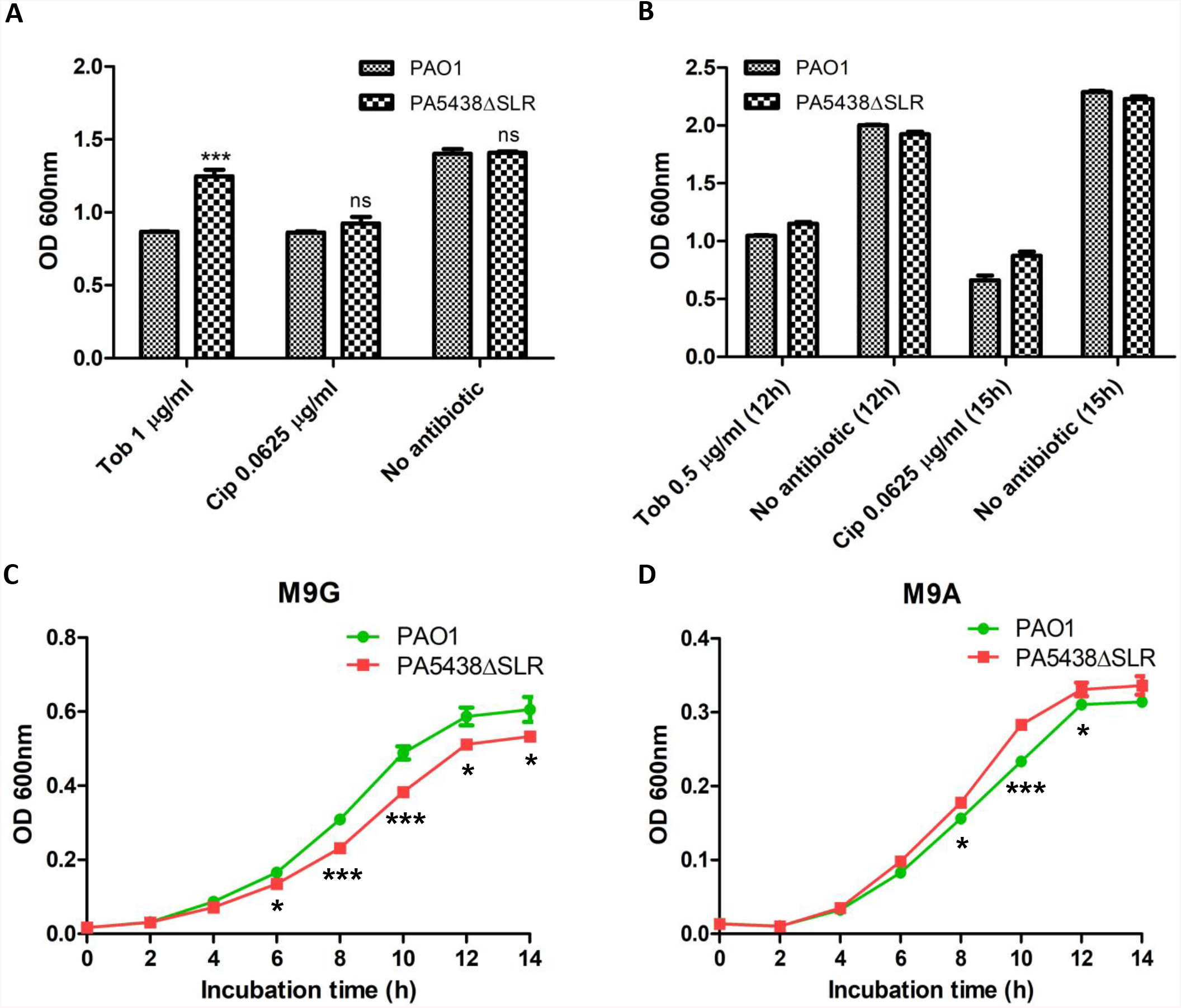
Antibiotic susceptibility test and comparison of growth under various conditions. **(A)** Antibiotic susceptibility tests with tobramycin (Tob) and ciprofloxacin (Cip) were performed. Initial CFU of PAO1 and the PA5438ΔSLR mutant were adjusted to 5 × 105 CFU, and OD_600nm_ was measured after 18 hours of static incubation in LB supplemented with 1 μg/ml tobramycin or 0.0625 μg/ml ciprofloxacin. Growth in the absence of antibiotics was measured as control. \*\*\**p*<0.001; ns: not significant **(B)** OD_600nm_ of PAO1 and the mutant after 12 hours of shaking incubation in LB with 0.5 μg/ml tobramycin and no antibiotic were measured. For OD_600nm_ measurement of the ciprofloxacin group, a final concentration of 0.0625 μg/ml was used and cultures were incubated for 15 hours with shaking. **(C)** Growth curves of PAO1 and the mutant in M9 minimal media supplemented with glucose as the sole carbon source were measured over 14 hours. \**p*<0.05; \*\*\**p*<0.001 **(D)** Growth curves of PAO1 and the mutant in M9 minimal media supplemented with acetate as the sole carbon source were measured over 14 hours. \**p*<0.05; \*\*\**p*<0.001

**Figure S3.**
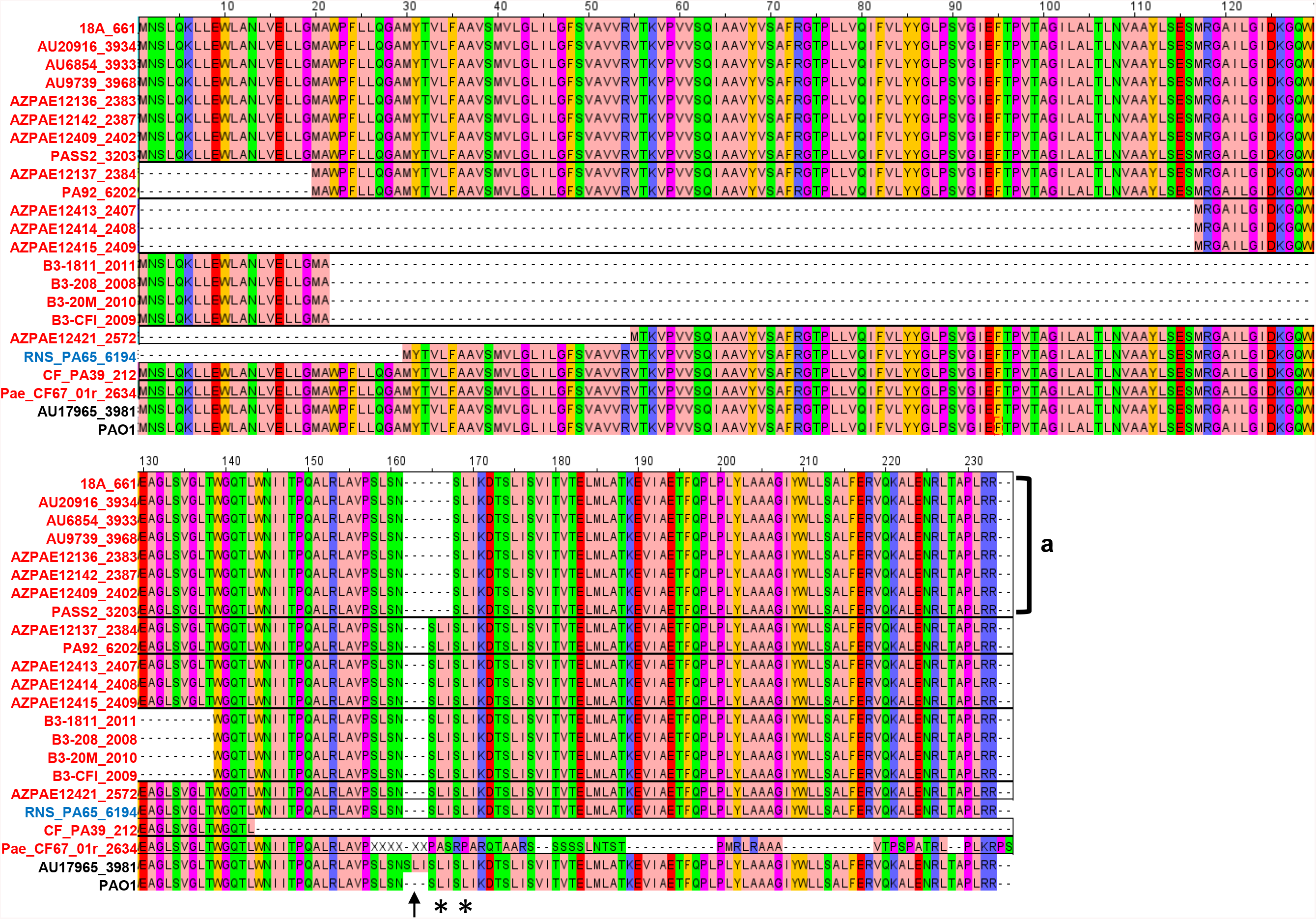
Multiple alignment of YecS and its homologues. Multiple alignment of YecS and its homologues is visually represented. Names of the genomes are shown to the left of the multiple alignment. PAO1 and AU17965_3981 are representative genomes of non-CF and CF groups, and genomes labelled by red and blue belong to the CF and non-CF groups, respectively. Black arrow marks where the additional SLI insertion occurs (162nd to 164th residues) compared to the YecS protein. Regions marked by asterisks are regions of SLI amino acid repeat sequences in YecS. Genomes in **a** contain a deletion of SLI, resulting in a single copy of SLI.

**Figure S4.**
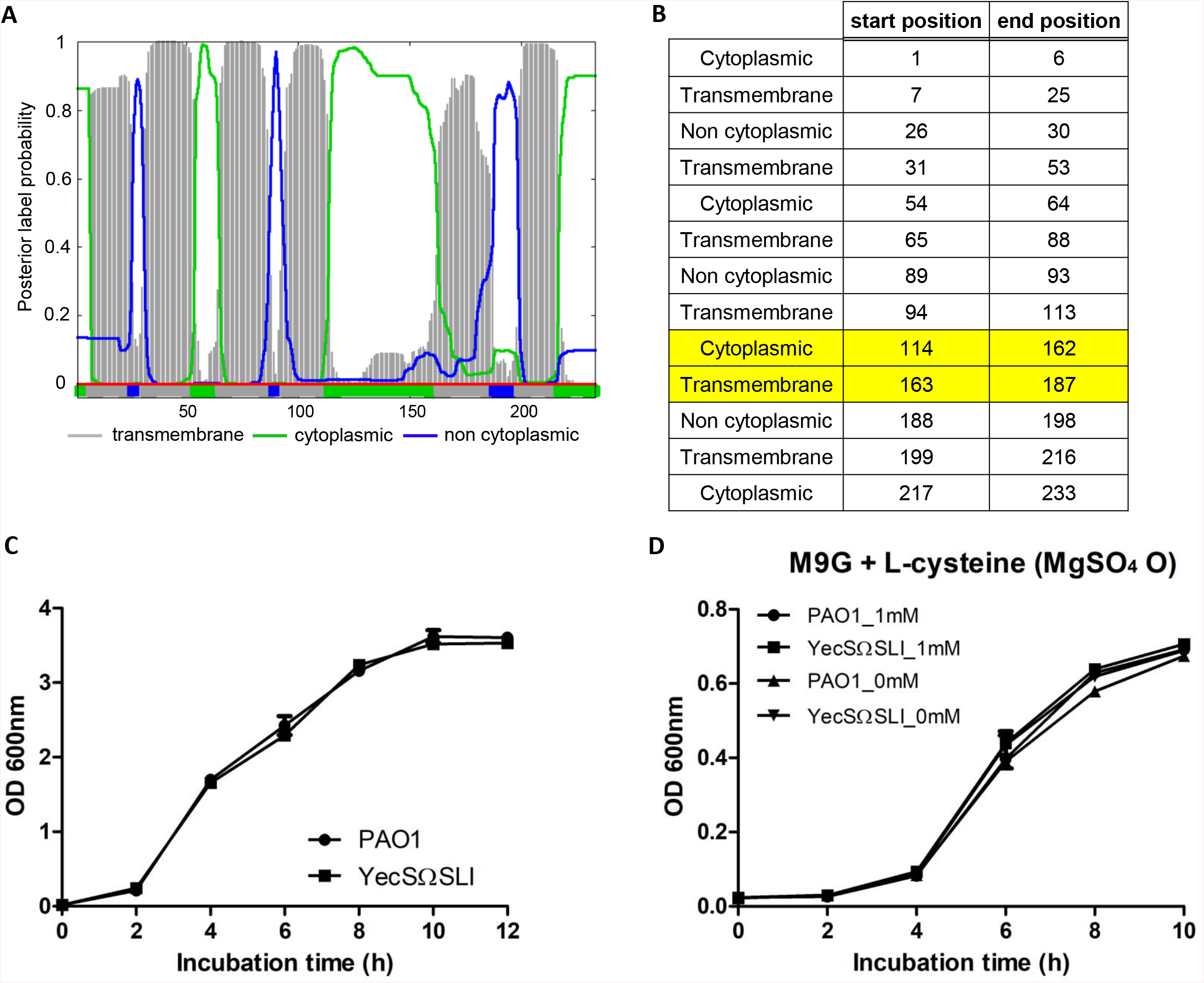
Transmembrane domain prediction and growth curves of the YecS mutant. **(A)** Predicted transmembrane domains of the AU17965_3981_04951 protein are portrayed. Numbers below the figure indicate the amino acid loci. **(B)** Detailed amino acid ranges of the predicted transmembrane domains are listed. The SLI insertion in the AU17965_3981_4951 protein is present within the region highlighted in yellow. **(C)** Growth curves of PAO1 and YecSΩSLI mutant in LB were measured over 12 hours. **(D)** Growth curves of PAO1 and YecSΩSLI in M9 minimal media supplemented with glucose, L-cysteine (1 mM) and MgSO_4_ (2 mM) were recorded over 10 hours.

**Table S1.** Blastp results of AceA and GlcB. Each columns represent: genomes (genomes containing SLR deletion in PA5438 homologues); pident (percentage of identical matches between reference gene and its homologue); length coverage (% of reference gene sequence covered by its homologue); Black boxes indicate non-CF genomes and red boxes indicate CF genomes. **(A)** Blastp results shown were conducted with the PAO1 AceA and its homologues from genomes containing SLR deletion in the PA5438 homologues. **(B)** Blastp results shown were conducted with the PAO1 GlcB and its homologues from genomes containing SLR deletion in the PA5438 homologues.

|  | genomes | pident(%) | Length coverage(%) |
| --- | --- | --- | --- |
|  | 14650_3305 | 99.812 | 100 |
|  | 18A_661 | 99.81 | 100 |
|  | AU17965_3981 | 99.812 | 100 |
|  | AU5471_3926 | 99.812 | 100 |
|  | AZPAE12137_2384 | 99.812 | 100 |
|  | AZPAE12149_2393 | 99.812 | 100 |
|  | AZPAE12153_2399 | 99.812 | 100 |
|  | AZPAE12409_2402 | 99.812 | 100 |
|  | AZPAE13757_2337 | 99.812 | 100 |
|  | AZPAE14816_2554 | 99.623 | 100 |
|  | COPD6d_6641 | 99.812 | 100 |
|  | DK2_174 | 99.812 | 100 |
|  | LES400_667 | 99.812 | 100 |
|  | LESlike4_669 | 99.812 | 100 |
|  | LESlike7_666 | 99.812 | 100 |
|  | PA102_7194 | 99.812 | 100 |
|  | PA59_6015 | 99.812 | 100 |
|  | PA66_5826 | 99.812 | 100 |
|  | Pae_CF67_01r_2634 | 99.812 | 100 |
|  | Pae_CF67_02a_2645 | 99.812 | 100 |
|  | Pae_CF67_05i_2710 | 99.812 | 100 |
|  | Pae_CF67_05p_2716 | 99.812 | 100 |
|  | Pae_CF67_06b_2829 | 99.812 | 100 |
|  | Pae_CF67_06c_2828 | 99.812 | 100 |
|  | Pae_CF67_06d_2830 | 99.812 | 100 |
|  | Pae_CF67_06e_2831 | 99.812 | 100 |
|  | Pae_CF67_06f_2832 | 99.812 | 100 |
|  | Pae_CF67_06g_2833 | 99.812 | 100 |
|  | Pae_CF67_06j_2720 | 99.812 | 100 |
|  | Pae_CF67_06l_2835 | 99.812 | 100 |
|  | Pae_CF67_06m_2836 | 99.812 | 100 |
|  | Pae_CF67_06n_2837 | 99.812 | 100 |
|  | Pae_CF67_06o_2847 | 99.812 | 100 |
|  | Pae_CF67_06p_2846 | 99.812 | 100 |
|  | Pae_CF67_06q_2838 | 99.812 | 100 |
|  | Pae_CF67_06r_2839 | 99.812 | 100 |
|  | Pae_CF67_06s_2840 | 99.812 | 100 |
|  | Pae_CF67_07p_2729 | 99.812 | 100 |
|  | Pae_CF67_08d_2732 | 99.812 | 100 |
|  | Pae_CF67_08f_2734 | 99.812 | 100 |
|  | Pae_CF67_08n_2742 | 99.812 | 100 |
|  | Pae_CF67_08q_2745 | 99.812 | 100 |
|  | Pae_CF67_08t_2748 | 99.812 | 100 |
|  | Pae_CF67_09l_2760 | 99.812 | 100 |
|  | Pae_CF67_10t_2787 | 99.812 | 100 |
|  | Pae_CF67_11c_2790 | 99.812 | 100 |
|  | Pae_CF67_12a_2799 | 99.812 | 100 |
|  | SCH_ABX04_5128 | 99.812 | 100 |

**Table S1. Blastp results of AceA and GlcB.**
|  | genomes | pident(%) | Length coverage(%) |
| --- | --- | --- | --- |
|  | 14650_3305 | 99.862 | 100 |
|  | 18A_661 | 99.72 | 100 |
|  | AU17965_3981 | 99.862 | 100 |
|  | AU5471_3926 | 99.862 | 100 |
|  | AZPAE12137_2384 | 99.862 | 100 |
|  | AZPAE12149_2393 | 99.862 | 100 |
|  | AZPAE12153_2399 | 99.862 | 100 |
|  | AZPAE12409_2402 | 99.862 | 100 |
|  | AZPAE13757_2337 | 99.862 | 100 |
|  | AZPAE14816_2554 | 99.724 | 100 |
|  | COPD6d_6641 | 100 | 100 |
|  | DK2_174 | 99.862 | 100 |
|  | LES400_667 | 99.862 | 100 |
|  | LESlike4_669 | 99.862 | 100 |
|  | LESlike7_666 | 99.862 | 100 |
|  | PA102_7194 | 99.862 | 100 |
|  | PA59_6015 | 100 | 100 |
|  | PA66_5826 | 100 | 100 |
|  | Pae_CF67_01r_2634 | 99.862 | 100 |
|  | Pae_CF67_02a_2645 | 99.862 | 100 |
|  | Pae_CF67_05i_2710 | 99.862 | 100 |
|  | Pae_CF67_05p_2716 | 99.862 | 100 |
|  | Pae_CF67_06b_2829 | 99.862 | 100 |
|  | Pae_CF67_06c_2828 | 99.862 | 100 |
|  | Pae_CF67_06d_2830 | 99.862 | 100 |
|  | Pae_CF67_06e_2831 | 99.862 | 100 |
|  | Pae_CF67_06f_2832 | 99.862 | 100 |
|  | Pae_CF67_06g_2833 | 99.862 | 100 |
|  | Pae_CF67_06j_2720 | 99.862 | 100 |
|  | Pae_CF67_06l_2835 | 99.862 | 100 |
|  | Pae_CF67_06m_2836 | 99.862 | 100 |
|  | Pae_CF67_06n_2837 | 99.862 | 100 |
|  | Pae_CF67_06o_2847 | 99.862 | 100 |
|  | Pae_CF67_06p_2846 | 99.862 | 100 |
|  | Pae_CF67_06q_2838 | 99.862 | 100 |
|  | Pae_CF67_06r_2839 | 99.862 | 100 |
|  | Pae_CF67_06s_2840 | 99.862 | 100 |
|  | Pae_CF67_07p_2729 | 99.862 | 100 |
|  | Pae_CF67_08d_2732 | 99.862 | 100 |
|  | Pae_CF67_08f_2734 | 99.862 | 100 |
|  | Pae_CF67_08n_2742 | 99.862 | 100 |
|  | Pae_CF67_08q_2745 | 99.862 | 100 |
|  | Pae_CF67_08t_2748 | 99.862 | 100 |
|  | Pae_CF67_09l_2760 | 99.862 | 100 |
|  | Pae_CF67_10t_2787 | 99.862 | 100 |
|  | Pae_CF67_11c_2790 | 99.862 | 100 |
|  | Pae_CF67_12a_2799 | 99.862 | 100 |
|  | SCH_ABX04_5128 | 99.862 | 100 |

CF genome non-CF genome

**Table S2.** Blastn results of promoter regions (upstream 200 bp) of *aceA* and *glcB*. Columns parallel those in Table S1. **(A)** Blastn results shown were conducted with the *aceA* promoter region in the PAO1 genome and its homologous promoters from genomes containing SLR deletion in the PA5438 homologues. **(B)** Blastn results shown were conducted with *glcB* promoter region in the PAO1 genome and its homologous promoters from genomes containing SLR deletion in the PA5438 homologues.

|  | genomes | pident(%) |
| --- | --- | --- |
|  | 14650_3305 | 100 |
|  | 18A_661 | 99.5 |
|  | AU17965_3981 | 100 |
|  | AU5471_3926 | 100 |
|  | AZPAE12137_2384 | 100 |
|  | AZPAE12149_2393 | 100 |
|  | AZPAE12153_2399 | 100 |
|  | AZPAE12409_2402 | 99.5 |
|  | AZPAE13757_2337 | 100 |
|  | AZPAE14816_2554 | 99.5 |
|  | COPD6d_6641 | 100 |
|  | DK2_174 | 100 |
|  | LES400_667 | 100 |
|  | LESlike4_669 | 100 |
|  | LESlike7_666 | 100 |
|  | PA102_7194 | 100 |
|  | PA59_6015 | 100 |
|  | PA66_5826 | 100 |
|  | Pae_CF67_01r_2634 | 100 |
|  | Pae_CF67_02a_2645 | 100 |
|  | Pae_CF67_05i_2710 | 100 |
|  | Pae_CF67_05p_2716 | 100 |
|  | Pae_CF67_06b_2829 | 100 |
|  | Pae_CF67_06c_2828 | 100 |
|  | Pae_CF67_06d_2830 | 100 |
|  | Pae_CF67_06e_2831 | 100 |
|  | Pae_CF67_06f_2832 | 100 |
|  | Pae_CF67_06g_2833 | 100 |
|  | Pae_CF67_06j_2720 | 100 |
|  | Pae_CF67_06l_2835 | 100 |
|  | Pae_CF67_06m_2836 | 100 |
|  | Pae_CF67_06n_2837 | 100 |
|  | Pae_CF67_06o_2847 | 100 |
|  | Pae_CF67_06p_2846 | 100 |
|  | Pae_CF67_06q_2838 | 100 |
|  | Pae_CF67_06r_2839 | 100 |
|  | Pae_CF67_06s_2840 | 100 |
|  | Pae_CF67_07p_2729 | 100 |
|  | Pae_CF67_08d_2732 | 100 |
|  | Pae_CF67_08f_2734 | 100 |
|  | Pae_CF67_08n_2742 | 100 |
|  | Pae_CF67_08q_2745 | 100 |
|  | Pae_CF67_08t_2748 | 100 |
|  | Pae_CF67_09l_2760 | 100 |
|  | Pae_CF67_10t_2787 | 100 |
|  | Pae_CF67_11c_2790 | 100 |
|  | Pae_CF67_12a_2799 | 100 |
|  | SCH_ABX04_5128 | 100 |

**Table S2. Blastn results of promoter regions (upstream 200 bp) of *aceA* and *glcB*.**
|  | genomes | pident(%) |
| --- | --- | --- |
|  | 14650_3305 | 100 |
|  | 18A_661 | 100 |
|  | AU17965_3981 | 99.5 |
|  | AU5471_3926 | 100 |
|  | AZPAE12137_2384 | 100 |
|  | AZPAE12149_2393 | 100 |
|  | AZPAE12153_2399 | 100 |
|  | AZPAE12409_2402 | 100 |
|  | AZPAE13757_2337 | 100 |
|  | AZPAE14816_2554 | 100 |
|  | COPD6d_6641 | 100 |
|  | DK2_174 | 100 |
|  | LES400_667 | 100 |
|  | LESlike4_669 | 100 |
|  | LESlike7_666 | 100 |
|  | PA102_7194 | 100 |
|  | PA59_6015 | 100 |
|  | PA66_5826 | 100 |
|  | Pae_CF67_01r_2634 | 100 |
|  | Pae_CF67_02a_2645 | 100 |
|  | Pae_CF67_05i_2710 | 100 |
|  | Pae_CF67_05p_2716 | 100 |
|  | Pae_CF67_06b_2829 | 100 |
|  | Pae_CF67_06c_2828 | 100 |
|  | Pae_CF67_06d_2830 | 100 |
|  | Pae_CF67_06e_2831 | 100 |
|  | Pae_CF67_06f_2832 | 100 |
|  | Pae_CF67_06g_2833 | 100 |
|  | Pae_CF67_06j_2720 | 100 |
|  | Pae_CF67_06l_2835 | 100 |
|  | Pae_CF67_06m_2836 | 100 |
|  | Pae_CF67_06n_2837 | 100 |
|  | Pae_CF67_06o_2847 | 100 |
|  | Pae_CF67_06p_2846 | 100 |
|  | Pae_CF67_06q_2838 | 100 |
|  | Pae_CF67_06r_2839 | 100 |
|  | Pae_CF67_06s_2840 | 100 |
|  | Pae_CF67_07p_2729 | 100 |
|  | Pae_CF67_08d_2732 | 100 |
|  | Pae_CF67_08f_2734 | 100 |
|  | Pae_CF67_08n_2742 | 100 |
|  | Pae_CF67_08q_2745 | 100 |
|  | Pae_CF67_08t_2748 | 100 |
|  | Pae_CF67_09l_2760 | 100 |
|  | Pae_CF67_10t_2787 | 100 |
|  | Pae_CF67_11c_2790 | 100 |
|  | Pae_CF67_12a_2799 | 100 |
|  | SCH_ABX04_5128 | 99.5 |

CF genome non-CF genome

**Table S3.**
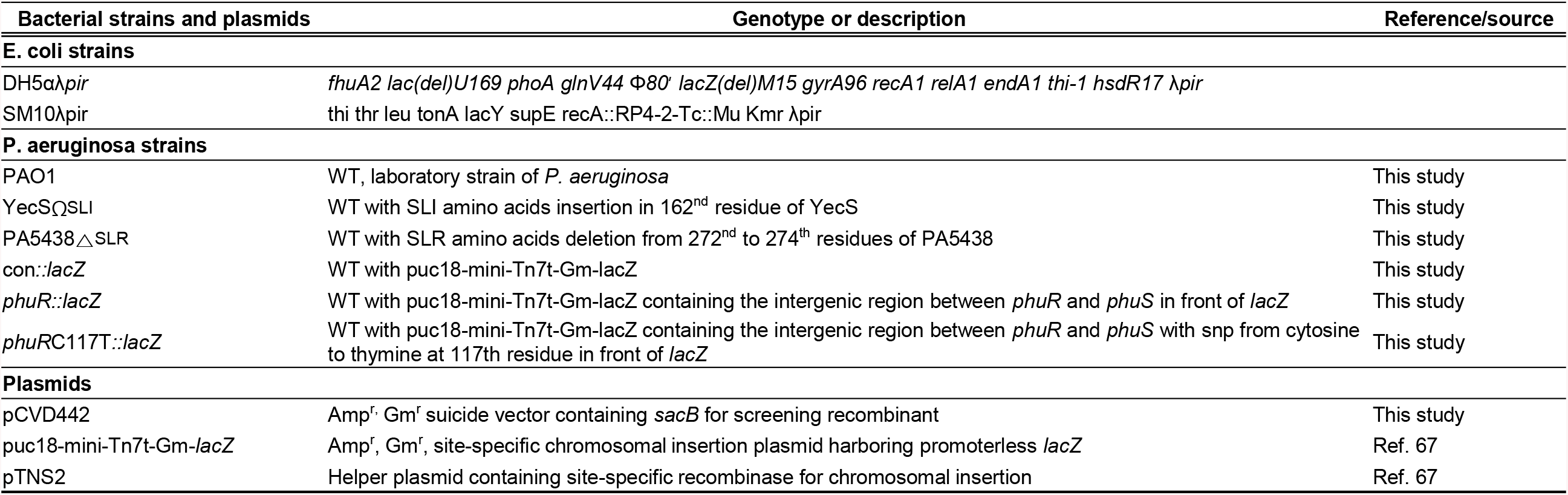
Strains and plasmids used in this study.

**Table S4.**
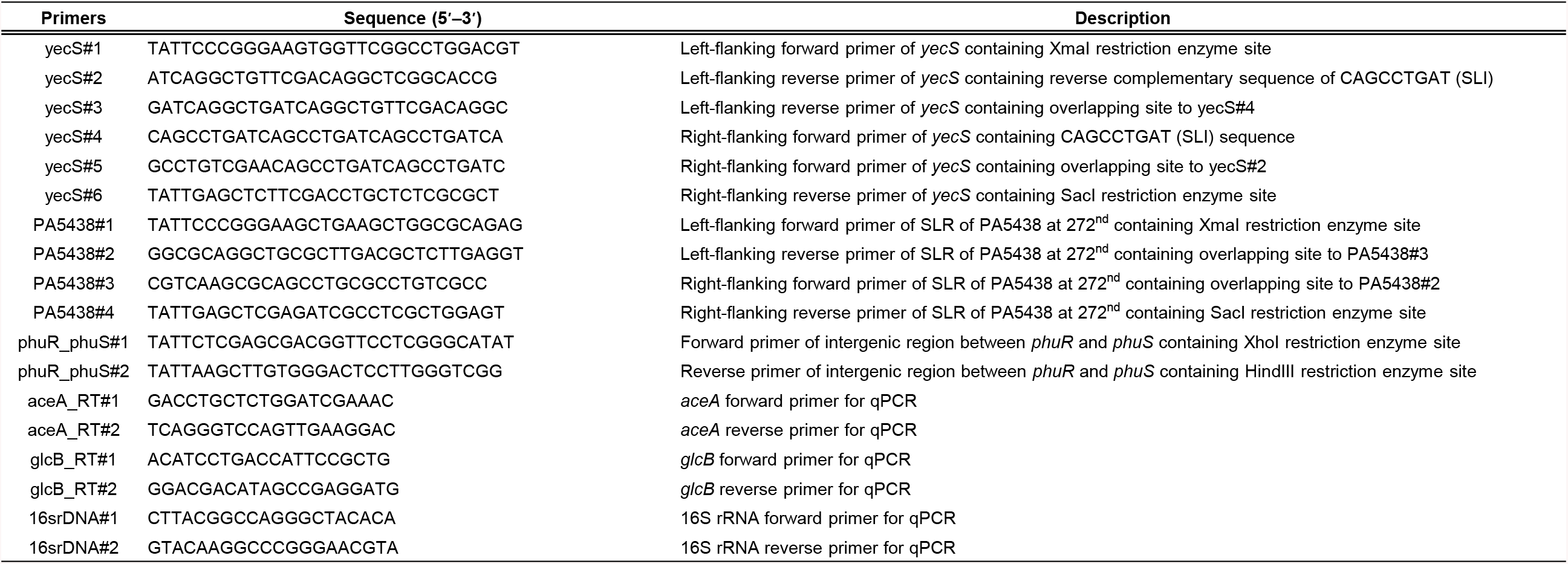
Primer information used in this study.

