## Supplementary Data 4 for "A Robust Genome-Wide Association Study Uncovers Signature Genetic Alterations among *Pseudomonas aeruginosa* Cystic Fibrosis Isolates"

**Amino acid sequences of candidate proteins in supplementary data 2, sheet 4.**

>105738_3985_01941 UDP-glucose 4-epimerase

MRVLVTGGAGFIGSHVLVELLGQGAKVVVLDNLVNGSSESLKRVERITGHPVGFVLGDIR

DSLLVERLLIDEKVDAVIHLAGLKAVGESVDDPLEYYESNVQGTISLLRAMQRVGVFKIV

FSSSATIYQMPGTLPISESSKVGGVASPYGRTKLTAEHMLDDLARSDARWSIAVLRYFNP

IGAHESGLIGEDPCGTPNNLLPYIAQVAVGRLSRLTVHGGDYPTIDGTGVRDYIHVCDLA

AGHTRALEYLRQGHGYHVWNLGTGTGYSVLQIIEAFERVSGRRIPFTVSGRRPGDVAECW

ADVSKAERELGWKAGLGLECMIADAWRWQVSNPSGYS

>AU10241_3928_00710 hypothetical protein

MRHWQRTIEQGNRCFVTGALIDAREHYLHALALAQVLLERWGDADEAVMAFVISHHNLAD

LHLQLEQPEETAEYLCACHERLLRVSADQKLPLALRQAAQHHSRRTYVELLSFIGEHGAY

PRTERLLKGPPNAEVRPEFLAPPPHFAITDGEFPCPTPCRPCLTPTMPWNRTSMR

>AU13210_3955_02428 Alginate biosynthesis protein AlgA

MNAVAPLIPCIVSGGSGTRLWPVSRESMPKPFMRLADDQSLLQKTFLRIAGLPDVARLLT

VTNRDLLFRTLDDYRAVNRSGLAQDLLLEPMGRNTAPAIAAAALHVQEHFGDQAQLLILP

ADHLIRDEQAFAAAVAEARGLAAQGYLVTFGITPERAETGFGYIEQGAPLGNGFRVARFV

EKPDQATAQSYLDSGKYLWNAGMFCFQAATVLQELERHAPEVLIAARAALADGSSLENGQ

CRQRELAAGAFAEAPDISVDYALMERSDKVAVVPCSIGWSDIGSWQALRELSAADENGNQ

VRGESVLHDVSNCYIDSPKRLVGAVGVHDLIIVDTPDALLVADAARSQDVKFVAQELKRR

GHDAFRLHRTVSRPWGTYTVLEEGRRFKIKRIVVRPKASLSLQMHHHRSEHWIVVSGMAL

VENGEREFLLNTNESTFIPAGHSHRLSNPGIIDLVMIEVQSGEYLGEDDIVRFNDIYGRA

PASDEKKA

>AU17965_3981_04951 L-cystine transport system permease protein YecS

MNSLQKLLEWLANLVELLGMAWPFLLQGAMYTVLFAAVSMVLGLILGFSVAVVRVTKVPV

VSQIAAVYVSAFRGTPLLVQIFVLYYGLPSVGIEFTPVTAGILALTLNVAAYLSESMRGA

ILGIDKGQWEAGLSVGLTWGQTLWNIITPQALRLAVPSLSNSLISLISLIKDTSLISVIT

VTELMLATKEVIAETFQPLPLYLAAAGIYWLLSALFERVQKALENRLTAPLRR

>AZPAE14712_2411_01394 Cell division protein ZapE

MTVVAVDGDQDHRLHPGRAEQRYWVVEAGQAGGFAELFARLSAGEVVSAQPIELAHRPLA

VRRHSESVLWCSYAQLCEVPLSALDFIGLCDRYRAILMDDLPCLSASQREGRIARGTEDG

VQLVEAGDRELPQLSVHDDGVRRFIALVDECYDRKVPLYLEARVPLEALYTEGYLAFAFR

RTLSRLREMQLARFGXAPRRRSGNCGWPPALIRRPGRGSVAVPAQQRVHVLRRRAALELQ

QFRGAVGEETVDAGRA

>AZPAE15072_2259_00011 hypothetical protein

MEFKTFRRTLLLAAVLFVPAGTSMAANLYAFGGRDIAVPSILPEGSVMSRYTFTPMQLCG

KPSCDLIGVSLYNKGSVWDPVDGPDLNTNVPGLSVRLLLDGIPASSRFKGNFSQIAEIQL

FRNSTPLSDGQFASGAFNSYFLITYKDGLIASGTSSIRLTGSVTTINATCQVADQTVKLQ

SIAAARLNGVGTYAAVTPFNLVVAGCPRGYNRVGYSLQAVGGAVAEGSGVLPRLAGSTAT

GVSIRVTDEAGVPLRMGLSLPVTAYDKNTGGAYWIPMKASYVQTAEKITPGSVLAAMVIL

LDYQ

>AZPAE15072_2259_00161 Quinone oxidoreductase 1

MAKRIQFAAYGGPEVLEYRDYQPAEPGPREVRVRNRAIGLNFIDTYYRSGLYPAPGLPSG

LGSEGAGEVEAVGSEVTRFKVGDRVAYATGPLGAYSELHVLAEEKLVHLPDGIDFEQAAA

VMLKGLTTQYLLRQTYELRGGETILFHAAAGGVGLFACQWAKALGVQLIGTVSSPEKARL

ARQHGAWETIDYSHENVARRVLELTDGKKCPVVYDSVGKDTWETSLDCVAPRGLLVSFGN

ASGPVTGVNLGILSQKGSLYVTRPTLGSYADTPEKLQAMADELFGLIERGDIRIEINQRF

ALAEAAKAHTELAARRTTGSTVLLP

>AZPAE15072_2259_00589 hypothetical protein

MNRIKHLLKDLAILIRANLWLIPVIAALVAAVFYFVAPPPPMSATMATGAEGGGYAVFAG

KLREKLKEQGFELKLVPSAGSRDNLERLLGTGEVDIALVQSGQERQLEAGQARQLQTLGA

VYQEPLWLFHRNNVHIDQLSDLLHLRLAIGSDGSGTRAATEAILQANDIAPGHYPITWEA

RGGNAVVDDLLAGKLDAAFIVGPAENPAIQKLADNDQLRLVNFRRSAAYEARLPFLKRVE

VGEGLLNLPRNVPERNISTLSPVATLVINERFHPALIPLVLETAREVMKDGSLLDKPGAF

PSAEPRTLRLHEDAERYYKSGLPLLQRYLPFRIASLADRYIILLIPFIAILIPLMKSIGP

LYRWRIRARIYRWYRYIRDIDRKLDSGTDAEQLRSEIERLEKLESELNTVEVPLSYYHEL

YELHLHLNFVIKRLHGLQERQATQAEAGNLG

>AZPAE15072_2259_00888 HTH-type transcriptional regulator BenM

MDLRQLRYFIAVAEELHFGRAAARLFISQPALSFDIKKLEEQLGTQLLLRNNKSVKLTGA

GQVLLVEARNLLLQAEKVKRLTQLSAEGDVGQLRVGFVNSMLYRGLPRAMSRFEREHPNM

EVVLGEMNSAEQAQALQRGQIDLGFVHWGRLPAEIVSEPLISDPFLCCLPAGHRLDGQAR

LDLAELRDEDFILFPRHVSPHYHDLIIARCVDAGFSPRIRHEARLWQTVAAMVGLGMGVA

LIPETLCLAWRNEVRYLEIEPAGARSEIHAILPASEPSRAAQAFLATLKSGLDDA

>AZPAE15072_2259_01307 hypothetical protein

MADVGSLGTRLLRIVLGLCALVLVLLALYVSLGRQLVPLVAEYRQQLEDEAGKQLGIPVR

IAELTGSWRGFEPLVVARDIQVGDGEQSLRLARVRLAPDLLGSLLARQVRIGSLELEGLK

LTLREGEDGQWSLDGLPHSDKPSDPRKLLQFLLQTQRISLLDSQLEVAPRGSAALSLSAV

GATLRSSSVGGQSLDARLQLPDGQPLALHAEGRIDSEDWPRSSARFYLSLPQSDWAQWLP

AGLTQEWKIVRAKAGGDFWFDWRDGKAQRLVARLLAPQLKASYAARKPVEINDLGMNLFF

DREAQGWKVRVGDLAANFGEQRWGEVELLLRRDQQNNEPHWKLQADRVDLTPLVPAIEAL

APLPDAAAEWVAGLKPKGILHNLNADFWPQREVPERVSYATNLEKVGISAFHEVPAVENV

SGTLTGTLAGGQLDASAQDFMLHLAKVFPEPWRYREARTRMFWSLDDRAFTLGSHLMRVE

GEEGRLAGDMLIRLMRDPGAEDYMDLQVGLSDGDARFTAKYLPTQLPGMNKSLANWLKTA

IRSGHVEQGYFQWQGSLNRGAVAEAHVMNLYFKVRDGELAYQPGWPALSKTVGEVFVEDS

GVRVLASSGNLLNSRVSDVKVDIPLGRPGQTPHLYVDGAVDSNLKDGIKLLQEAPIPTRK

IFAGWEGDGPLQGHLKLDIPLDHDEAGKTGVVVDFSTVGATLKMPSPKLDMSEVKGDFRF

DLAKGLSAPAVQARVLGSEVRGRIVAEGRGDARTRLLLNGQVAVKSLSDWLGAGQRPLPV

SGRLPFQLNLLLDGKDSQLQIDSDLKGAVVDLPAPFGKTAAQARPTQWRMTLDGAERRYW

ARYDGLASLAYAAPADKPLNGRGALRLGGDPALLPSAQGLRVRGRLAELDWDAWQATLKR

YGNGDQAASSAAGLLRGADLRIDSFKGFGQELKNLTVDLARQERAWQLVLVSDLASGRLV

LPDARGAPIVVDLDRLNLPKSTLPDENKVEDSDPLAAVDPRSLPAVDVKIGQVALGGQPL

GAWSLKVRPGSNGVAFNDLDLDLRGLHVNGSLRWDGSPGNTRSTYQGRIEGKNLADVLKA

WNFAPSATSERFRMDINGQWPGSPAYMALKRFSGSMDASLRKGQFVEVEGSAQALRVFGL

LNFNAINRRLRLDFSDILGKGLSYDRVKGGLSATDGVYVTREPLKLEGPSSNLELNGTLD

LAHDRIDAKLLVTLPLTNNLTLAALIVGGPAVGGAVFVVDKLLGDRVSRFASVQYSVKGP

WQDPKISFDKPFEKPR

>AZPAE15072_2259_02084 hypothetical protein

MQRSIATVSLSGTLPEKLEAIAAAGFDGVEIFENDLLHYDGSPRDVRRLCADLDLEILLF

QPFRDFEGCRRERLERNLERAERKFDLMQELGTDLVLVCSNVAADALGEPALLADDLRQL

AERAAVRGLRIGYEALAWGRQVNTWEQAWDLVRRADQANLGLILDSFHTLSLDGDPRGIA

DLPGEKIFFVQMADAPLLAMDVLEWSRHFRCFPGQGGFDLAGFLAPVVASGYRGPLSLEV

FNDGFRAAPTRANAVDGLRSLLYLEEKTREHLQRQTPHVAVDELFAPPPASLCDGIEFLE

FAVDKTLSARLGQWLQRLGFARAGEHRSKNVSLLRQGDINLVLNAEPYSFAHGFFEAHGP

SLCATALRVRDAGQALERARAYGGQPYRGLLGPNEREIPAVRALDGSLLYLVERHTEGRS

IYDSDFVTNDADTSGLGLRRVDHVALALPAEGLDSWVLFYKSLFDFGADDEVVLPDPYGL

VTSRAVRSPCGSVRLPLNISEDRNTAIARSLSSYRGSGVHHIAFDCADIFAAVAQAKEAG

VALLEIPLNYYDDLAARFDFDDEFLSELAYYNVLYDRDAQGGELFHVFTEPFEERFFFEI

LQRRHGYAGYGAANVPVRLAAMAQARRGVRRVKL

>AZPAE15072_2259_02833 hypothetical protein

MSQSLKRALFSALLVILVSYPILGLKLRTVGIKLEVLGADAQTLWTIAAAALAMFVWQLF

RDRIPLKLGRGVGYKVNGSGLKNFLSLPSTQRWAVLALVVVAFVWPFFASRGAVDIATLI

LIYVMLGIGLNIVVGLAGLLDLGYVGFYAVGAYTYALLAEYAGFGFWTALPIAGMMAALF

GFLLGFPVLRLRGDYLAIVTLGFGEIIRILLRNMTEITGGPNGIGSIPKPTLFGLTFERR

AQEGMQTFHEFFGIAYNTNYKVILLYVVALLLVLLALFVINRLMRMPIGRAWEALREDEV

ACRALGLNPTIVKLSAFTIGASFAGFAGSFFAARQGLVTPESFTFIESAMILAIVVLGGM

GSQLGVILAAVVMVLLQEMRGFNEYRMLIFGLTMIVMMIWRPQGLLPMQRPHLELKP

>AZPAE15072_2259_03078 Multidrug resistance protein Stp

MNPSRRVALVAAIYLGTFIASLDISIVNLALPTLQYALDTDLAGLQWVVDAYALCLSAFM

LSSGPLSDRYGRKLTWLLGVGLFSFGSLLCALATSLPLLLFGRAVQGIAGALLIPGALSI

LTQAFHDPGQRAQVIGGWTSFSALSLILGPLLGGLLVEHAGWQSIFLINLPLGLLALALG

LWGIEETAHPEHAAFDPLGQLLSVVWLGALTYALIAAGESGWLSPTAWPALLLAGVGLLG

FLFVERRTARPLLPLGLFRQAGFAVCNLASFVLGFSGYASLFFLSLFFQQVQGASAQQAG

FYLAPQFLAMGALSMLFGRLQRHVPLRRLLVLGYLVIGLAMLALAACGTGTAYPWVGLLL

VALGLGMGLAVPGTGLAVMASVARERSGMASATMNTLRQAGMAVGIALLGALLSGRAIVV

LGDRLEELGIADAQRLATQAITAHRLPGSLAGLDAELPAALAEGFRLAMLVAGASALLAA

ALLWRLRVSAGPAADTVGASGRTAGVQLQADRR

>AZPAE15072_2259_03267 4-hydroxyphenylacetate 3-monooxygenase oxygenase component

MKPEDFRASATRPFTGEEYLASLRDDREIYIYGDRVKDVTSHPAFRNAAASMARLYDALH

DPQSKEKLCWETDTGNGGYTHKFFRYARSADELRQQRDAIAEWSRLTYGWMGRTPDYKAA

FGSALGANPGFYGRFEDNAKTWYKRIQEACLYLNHAIVNPPIDRDKPVDQVKDVFISVDE

EVDGGIVVSGAKVVATNSALTHYNFVGQGSAQLLGNNTDFALMFIAPMNTPGMKLICRPS

YELVAGIAGSPFDYPLSSRFDENDAILVMDKVFIPWENVLIYRDFERCKQWFPQGGFGRL

FPMQGCTRLAVKLDFITGALYKALQCTGSLEFRGVQAQVGEVVAWRNLFWSLTDAMYGNA

SEWHSGAFLPSAEALQAYRVLAPQAYPEIKKTIEQVVASGLIYLPSGVRDLHNPQLDKYL

STYCRGSGGMGHRERIKILKLLWDAIGSEFGGRHELYEINYAGSQDEIRMQALRQAVGSG

AMKGMLGMVEQCMGDYDENGWTVPHLHNPDDINVLDRIRQ

>AZPAE15072_2259_03600 hypothetical protein

MLELTNAQIGGWIASFVLPLFRVAALLMTMPVIGTQLVPVRVRLYLALGVCVVLVPNLPP

MPQVDALSMKAMLLIGEQILVGALLGFSLQLLFHAFVIAGQIISMQMGLGFASMVDPANG

VSVPVLGQFFTMLVTLLFLAMNGHLVVFEVIAESFVTLPVGEGLSGNHFWIIAGKLGWVM

GAALLLALPAITALLVVNLAFGAMTRAAPQLNIFSIGFPLTLVLGLVILWIGTADLLSQY

QLLAGEALQFLRELVRAK

>AZPAE15072_2259_04340 D-alanyl-D-alanine carboxypeptidase DacB

MFKSLRTLAFATLLPFALPTLAQVNATLPANVQKALQTNKLTGNDLSLVLIPLDGPGNPT

YYNADVSVNPASTMKLFTTYAALEMLGPTYQWKTEFYTDGQLKNGVLNGNLYLKGGGDPK

LNMEKLWLLMRDLRANGVTKVTGDLVLDRSYFNIPQLPVFNDDGGDDTKPFLVGPDSLLV

NLKSVRMVVRTDGNKVNVQMDPPLANVRIDNQVKMTAPATCPAWPKLRFSPVTQFDGTTL

LATGQIPQGCSAQTYMSLLDHPGYTAGAVRGIWQELGGSILGKDRQGSVPRNATLIAKAF

SPDLVEIIRDINKYSNNTMARQLFLSIGAQFRNSADGDDAQAAQRVVRQWLARKGITVPR

LVMENGSGLSRQERVSAREMAAMLQAAWHSPYAPEYISSLPLAGLDGTMRKRLRRTALVG

EAHVKTGTLNTVRALAGFSRDASGHNWVVVAILNSPRPWGASAILDQVLLSLHARK

>AZPAE15072_2259_05171 hypothetical protein

MPNFGFHIAPAHPVAGRLIYDSKKLAENILKQHSDERVFSRAQEQKRLSEGDVVGGAPCC

KAIHITLGFDGTNNNDKADGSSVSPSCSNVARLIHASIGSGDDINSRGIFKYYCPGVGTV

FPDIKEFTPSNMGLIGAEGGENRINWGLVQLVDALFYTLLKSRLKLNEVQGLVEEMSTNW

TVSTLTGGLLENGEKKRRTALEPKLKELEEKLRQRQNSGQKPHILAMRLYKHEPLPTGCR

S
